# An empirical three-dimensional metric field for color space

**DOI:** 10.64898/2026.03.09.710376

**Authors:** Jan Koenderink, Andrea van Doorn, Doris I. Braun, Karl R. Gegenfurtner

## Abstract

A complete empirical characterization of color discrimination in three dimensions has long remained out of reach. Classical studies, beginning with MacAdam’s ellipses, provided local measurements in restricted chromatic planes, but a spatially dense and internally consistent mapping of discrimination structure across full color space has not yet been achieved. Here we present such a systematic three-dimensional measurement of color discrimination in the standardized sRGB stimulus space.

Eight observers measured discrimination regions at 35 reference colors distributed on a body-centered cubic lattice within the RGB cube. At each location, color differences were probed along seven orientations, yielding 14 directional extents. These measurements defined centrally symmetric convex regions that were fitted with minimum-volume ellipsoids, providing a compact description of local discrimination structure. Ellipsoids were represented as symmetric positive-definite matrices and analyzed using a Frobenius geometry, enabling normalization across observers and smooth interpolation to arbitrary locations.

The resulting metric field is spatially smooth, highly structured, and remarkably consistent across observers up to an individual global scale factor. Grain size increases along the achromatic axis and exhibits systematic chromatic asymmetries. Comparison with CIEDE2000 reveals substantial agreement in overall scale variation but systematic differences in local anisotropy. Because the metric field is defined geometrically, it can be transformed into linear RGB or any other color representation through the corresponding coordinate transformations.

Together, these data provide a coherent three-dimensional empirical mapping of color discrimination across a large ecologically relevant volume of color space and establish an empirical framework for perceptual color metrics.

## Introduction

Color vision was among the earliest domains of vision science to receive a rigorous mathematical treatment. Since the nineteenth century, its foundations have been shaped by the principles of trichromacy and univariance, which together imply that color matches can be represented in a linear three-dimensional vector space (Young, 1802; Helmholtz, 1867; Maxwell, 1857; Grassmann, 1853; Schrödinger, 1925; Krantz, 1975). These principles have proven to be extraordinarily successful. They account for color matching and a wide range of perceptual phenomena and provide the conceptual and computational basis for virtually all color measurement and reproduction, from displays and print media to painting and digital imaging (e.g., Judd, 1953; Wyszecki & Stiles, 1982; Koenderink, 2010; Berns, 2016). Yet despite this long history and firm formal foundation, color space lacks a generally accepted perceptual metric. Although colors can be represented as points in a three-dimensional vector space, separations in this space do not correspond in any simple or uniform way to perceived color differences. The problem of defining a perceptually meaningful metric has therefore remained central to color science from its inception. For a recent historical review, see Mollon and Danilova (2026).

The absence of a natural subjective metric was recognized early. After formalizing trichromacy, Helmholtz (1896) proposed a Riemannian metric for color space based on Weber’s law. Schrödinger (1920) proposed a modification aligned with the emerging luminous efficiency function V(λ). Several related line-element approaches were subsequently proposed, but none survived later experimental scrutiny, as they failed to accommodate increasingly extensive psychophysical data.

A decisive shift occurred when the problem was approached empirically rather than theoretically. The most influential contribution in this direction is MacAdam’s classical study of chromatic discrimination ellipses (MacAdam, 1942). MacAdam measured thresholds in an isoluminant plane. His ingenious apparatus allowed observers to adjust mixtures of primary lights along differently oriented lines, yielding distributions of matches that were well approximated by ellipses. His findings revealed that perceptual color space is locally anisotropic and spatially inhomogeneous. Subsequent work extended these measurements into three dimensions at selected chromaticity locations. Brown and MacAdam (1949) reported discrimination ellipsoids elongated predominantly along the luminance axis. Wyszecki and Fielder (1971), measuring additional observers and repeating measurements within observers, documented considerable irregularity and, importantly, substantial variability across repeated sessions of the same observer. Together, these studies underscored both the geometric complexity of perceptual color space and the experimental fragility of mapping it comprehensively.

### The curse of dimensionality

The time and effort required for these classical threshold measurements appear to have discouraged systematic attempts to map discrimination structure throughout three-dimensional color space. Any such attempt immediately encounters a fundamental obstacle: the curse of dimensionality (Bellman, 1957).

Psychophysical discrimination thresholds are typically on the order of a few percent along a single dimension. If one were to sample RGB space with even modest resolution—say, 10 distinguishable steps per axis—this would imply 10^3^ (1,000) distinct reference locations. At each location, discrimination would in principle need to be probed along multiple directions. Even the coarsest directional sampling along 6 directions per location multiplies the required measurements substantially, requiring about 6,000 psychometric function estimates. Exhaustive empirical mapping therefore becomes combinatorially infeasible using conventional paradigms.

This combinatorial burden has strongly shaped the history of color metrics. It has favored sparse local measurements, strong parametric assumptions, and extensive interpolation. Very specific questions were answered for relatively small regions of color space (e.g., Wandell, 1985; Poirson & Wandell, 1990; Krauskopf & Gegenfurtner, 1992; Witzel & Gegenfurtner, 2015; Danilova & Mollon, 2016; Hedjar, Toscani & Gegenfurtner, 2025a, 2025b), including geometric analyses of local color-space structure (Wuerger, Maloney & Krauskopf, 1995; Ennis & Zaidi, 2019), while only a few studies have attempted systematic exploration of color geometry across larger regions of space (Griffin & Mylonas, 2019; Koenderink, van Doorn & Gegenfurtner, 2018; Hong et al., 2025).

As a result, color science faces a paradox: although color is among the most intensively studied sensory domains, the empirical basis for global perceptual distance relations across full color space remains remarkably limited.

### The current standard

Following MacAdam’s measurements of chromaticity discrimination ellipses, numerous efforts sought transformations that would render these ellipses approximately circular, enabling a simple Euclidean representation of color distances (e.g., Hunter, 1942). In parallel, perceptual scaling of the Munsell color system aimed to achieve approximately uniform spacing of color samples (Hunter, 1952; Nickerson, 1950). These strands converged in the CIE 1976 uniform color space CIELAB and the associated color-difference measure ΔE*₇₆* (see Fairchild, 1998; Wyszecki & Stiles, 1982). The current standard, CIEDE2000 (Luo, Cui, & Rigg, 2001; see Kuehni, 2003), represents decades of empirical refinement of ΔE*₇₆*. Based on accumulated suprathreshold discrimination datasets (e.g., BFD, RIT–DuPont, Leeds, Witt), it introduced lightness, chroma, and hue weighting functions, as well as a rotation term to account for systematic deviations in the blue region. These developments significantly improved industrial tolerance prediction (e.g., Luo et al., 2022).

However, CIEDE2000 does not define a globally consistent perceptual metric. Rather, it is a locally parameterized color difference formula. Its structure presupposes separable lightness, chroma, and hue components, and the empirical data on which it rests are comparatively sparse and largely confined to planar sections of color space. A complete three-dimensional empirical characterization of perceptual color geometry is still lacking, despite recent progress in two dimensions (Hong et al., 2025). We address this gap by measuring a three-dimensional empirical metric field throughout the standardized sRGB stimulus space.

### Why RGB matters

Two independent considerations make the standardized sRGB stimulus space a particularly suitable domain for such a measurement.

First, RGB is the generative space of modern visual technology. Contemporary displays such as LCD, OLED, LED projection, and immersive head-mounted systems produce color through additive mixtures of three primaries. Any psychophysical measurement performed on a display is therefore inherently embedded in RGB coordinates. Measuring the perceptual metric directly in this space avoids the additional transformations and assumptions that would be required if stimuli were first mapped into a derived color representation. Moreover, the use of standardized sRGB primaries facilitates transformation of both stimuli and metric fields into other coordinate systems such as linear RGB, XYZ, LMS, CIELAB or DKL space.

Second, RGB space is of great ecological validity. Contrary to conventional wisdom based on two-dimensional chromaticity diagrams, the RGB cube occupies a substantial fraction of the physically realizable color solid and in fact the largest such fraction attainable (Koenderink, van Doorn & Gegenfurtner, 2021). When projected into CIE xy space, the RGB gamut appears as a small triangle, surrounded by a larger region, because the projection dramatically distorts volumetric relations. In three dimensions, the RGB cube exhausts about two-thirds of the volume of the color solid, and it is actually quite hard to find spectra of real objects that are situated in the area outside of the triangle. Figure 1 illustrates this point. Most naturally occurring surface reflectances fall comfortably within the sRGB gamut, and only a small fraction of Munsell samples lie outside it. Only certain highly saturated self-luminous stimuli, including signal lights and instrument panels, extend beyond the sRGB gamut.

**Figure 1:**
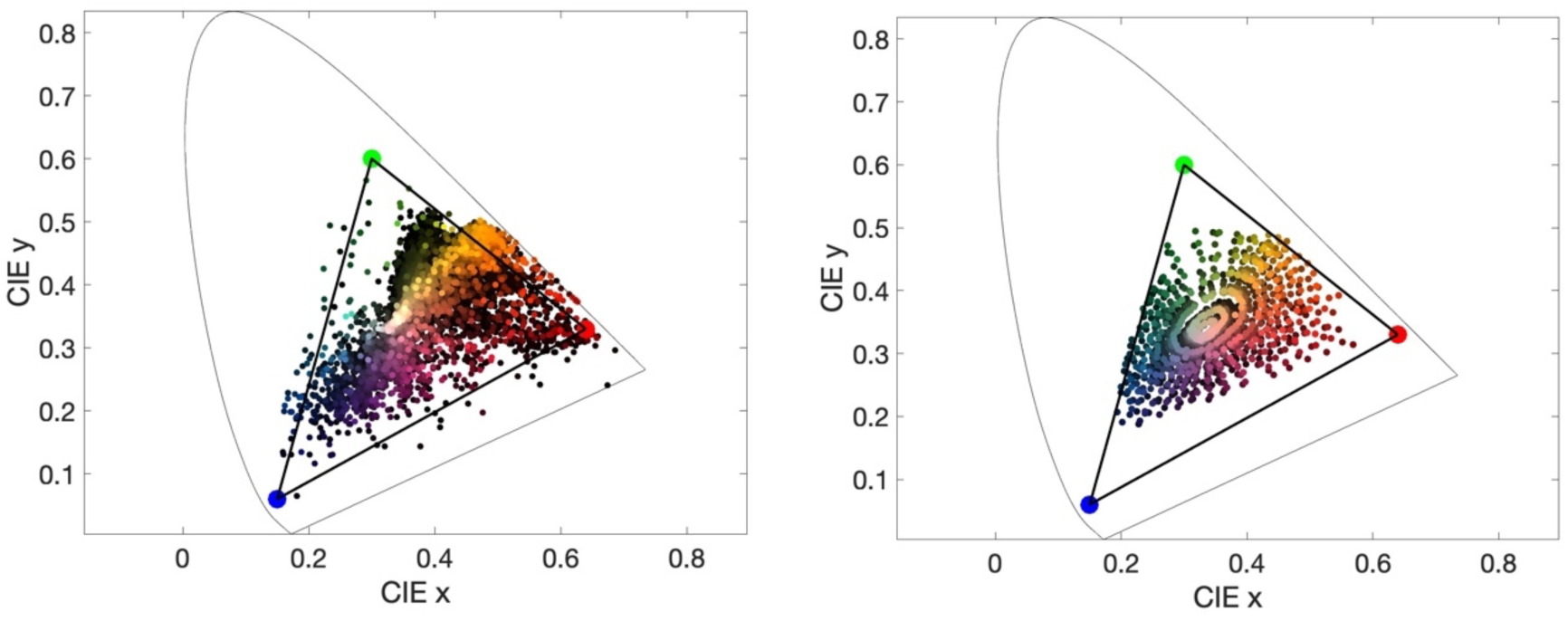
Spectral distributions in the CIE xy chromaticity diagram. The triangle indicates the sRGB color gamut. In the left panel, 6474 natural reflectances are plotted from several databases (see Akbarinia & Gegenfurtner, 2018). In the right panel, Munsell chips are plotted.

Thus, RGB space is not merely a technological convenience. It captures the dominant range of ecologically encountered surface colors and provides a physically generative three-dimensional stimulus space suitable for volumetric sampling. This makes it a principled domain in which to measure a complete three-dimensional perceptual metric field.

In the present work, measurements were performed directly in gamma-encoded sRGB coordinates. This choice reflects the native representation of modern display systems and provides a convenient, approximately perceptual scaling of intensity, analogous to the use of decibels in acoustics (see Appendix A1). Importantly, the resulting metric field is not tied to sRGB. Transformations to linear RGB are associated with only minor geometric changes (Appendix A3). Once represented in linear RGB, the field can be transformed exactly into XYZ, LMS, DKL, and other linearly related color spaces.

### The metric field

Rather than seeking a single global distance formula, we approach the problem at a more fundamental level. We treat color space as endowed with a metric field, a spatially varying local structure that specifies, at each color, which neighboring colors are perceptually similar and which are notably different. Operationally, this metric field is characterized by measuring, around each reference color, the fuzzy region of colors that are not qualitatively distinguishable from it. We assume that these regions are convex and centrally symmetric, with boundaries that are well approximated by ellipsoids. We further assume that these ellipsoids deform smoothly as the reference color varies. Under these conditions, the field can be reconstructed from a finite set of local samples, with interpolation filling occasional lacunae according to local rates of change.

This perspective shifts the focus from isolated discrimination thresholds to the global organization of perceptual color geometry. Instead of asking how small a difference can be detected at a single location, we ask how local regions of qualitative equivalence are arranged throughout a three dimensional stimulus space. Such regions have a long history in color science, from the color atlases of Munsell (1912) and Ostwald (1919) to Wright’s (1941) colorimetric studies, and more recent studies (Boynton & Kambe, 1980; Griffin & Mylonas, 2019; Koenderink, van Doorn & Gegenfurtner, 2018).

Our primary goal in this study is to obtain these fundamental empirical measurements across RGB space. Once established, the metric field specifies perceptual color differences by interpolation at any location and in any direction. The field is deliberately independent of any particular theory of neural processing. Instead, it provides a comprehensive three-dimensional empirical reference against which neural models, computational theories, and existing color-difference formulae can be quantitatively evaluated.

## Methods

In the present study, observers judged notable qualitative differences rather than just noticeable differences. For each reference color, we estimated the region of neighboring colors that were not yet experienced as qualitatively distinct. These regions were subsequently approximated by ellipsoids and interpreted as local descriptors of the perceptual metric field introduced above.

All measurements were performed directly within the RGB cube. As justified above, RGB defines the generative stimulus space of the experiment. Observers adjusted color differences along specified directions in RGB coordinates. No transformation into CIE or cone excitation spaces was applied during measurement. The goal was to estimate perceptual structure in the native three dimensional stimulus domain in which the sampling is defined.

This objective motivated several methodological choices.

Observers did not perform forced-choice judgments (“same vs. different”). Instead, they adjusted color differences until a clear qualitative distinction was experienced. Although such judgments are necessarily criterion-dependent, global differences in criterion level can later be normalized by scale, leaving spatial structure intact.

Color differences were measured using a method of adjustment with reciprocal bracketing. Along predefined directions in RGB space, observers increased deviation from a reference color until a qualitative step was perceived, then reversed direction and bracketed the transition repeatedly. Multiple (5) turning points were recorded, and medians were used to estimate directional extents.

Stimulus geometry was designed to eliminate edge-detection strategies. Colors were presented as concentric circular regions separated by a thick black outline, preventing reliance on chromatic edges. Judgments were therefore based on area color rather than boundary contrast.

Finally, chromatic adaptation was controlled by embedding the stimuli in a heterogeneous mosaic background sampling the RGB cube. This maintained the perceptual domain of related colors and reduced adaptation to any specific reference. As with any psychophysical measurement of color discrimination, the resulting metric field is therefore defined relative to the adaptation state induced by the stimulus configuration rather than independently of viewing conditions.

Together, these procedures define an operational method for estimating local regions of qualitative color equivalence in RGB space.

### Observers

Eight observers participated in the main experiment. In addition to the four authors (all aged 64 years and older), four young lab members took part who were naïve with respect to the purpose of the study. All reported normal color vision and normal or corrected-to-normal visual acuity. Participants viewed the display binocularly in a darkened room at a comfortable reading distance. If necessary, they wore their habitual optical correction. Each observer completed five sessions of approximately one hour each. The study was approved by the local ethics committee (Ethics approval # LEK 2020-0015) and conducted in accordance with the Declaration of Helsinki.

### Stimuli and Sampling

Stimuli were presented on a calibrated Apple MacBook Liquid Retina XDR display operating under standard sRGB intensity encoding. The display was characterised with a Konica Minolta CA-2000. The measured CIE xyY coordinates of the primaries at full drive were red x = 0.639, y = 0.334, Y = 22.9 cd/m²; green x = 0.310, y = 0.602, Y = 80.0 cd/m²; and blue x = 0.142, y = 0.074, Y = 8.3 cd/m². The white point was x = 0.306, y = 0.320, with a peak white luminance of 112 cd/m² and a black level of 0.4 cd/m², and the resulting gamut adhered to sRGB within measurement accuracy. Measured display gamma values were 2.47 for red, 2.42 for green, and 2.42 for blue, consistent with the sRGB transfer function. The three channels were additive, their summed spectra reconstructing the white spectrum to within 1%. The characteristics of our display are provided in Appendix A5.

All experiments were performed directly in gamma-encoded sRGB coordinates rather than after linearization. This choice preserves the native display representation and yields a compressed intensity axis qualitatively similar to the nonlinear lightness encoding used in perceptually motivated spaces such as CIELAB L* (see Appendix A1). Transformations between nonlinear and linear RGB are smooth and invertible, allowing the resulting metric field to be mapped into linear RGB, LMS, XYZ, or related spaces (see Appendix A3).

Each trial consisted of a 5 degree central disk (adjustable comparison color) surrounded by a 10 degree annulus (reference color). Figure 2A shows an example stimulus display. The regions were separated by a thick black 4 min arc wide circular outline to eliminate edge-based cues. Stimuli were embedded in a dynamic Voronoi mosaic background sampling the distribution of naturally occuring object colors within the RGB cube. The heterogeneous background maintained a stable domain of related colors and reduced chromatic adaptation to the reference.

**Figure 2:**
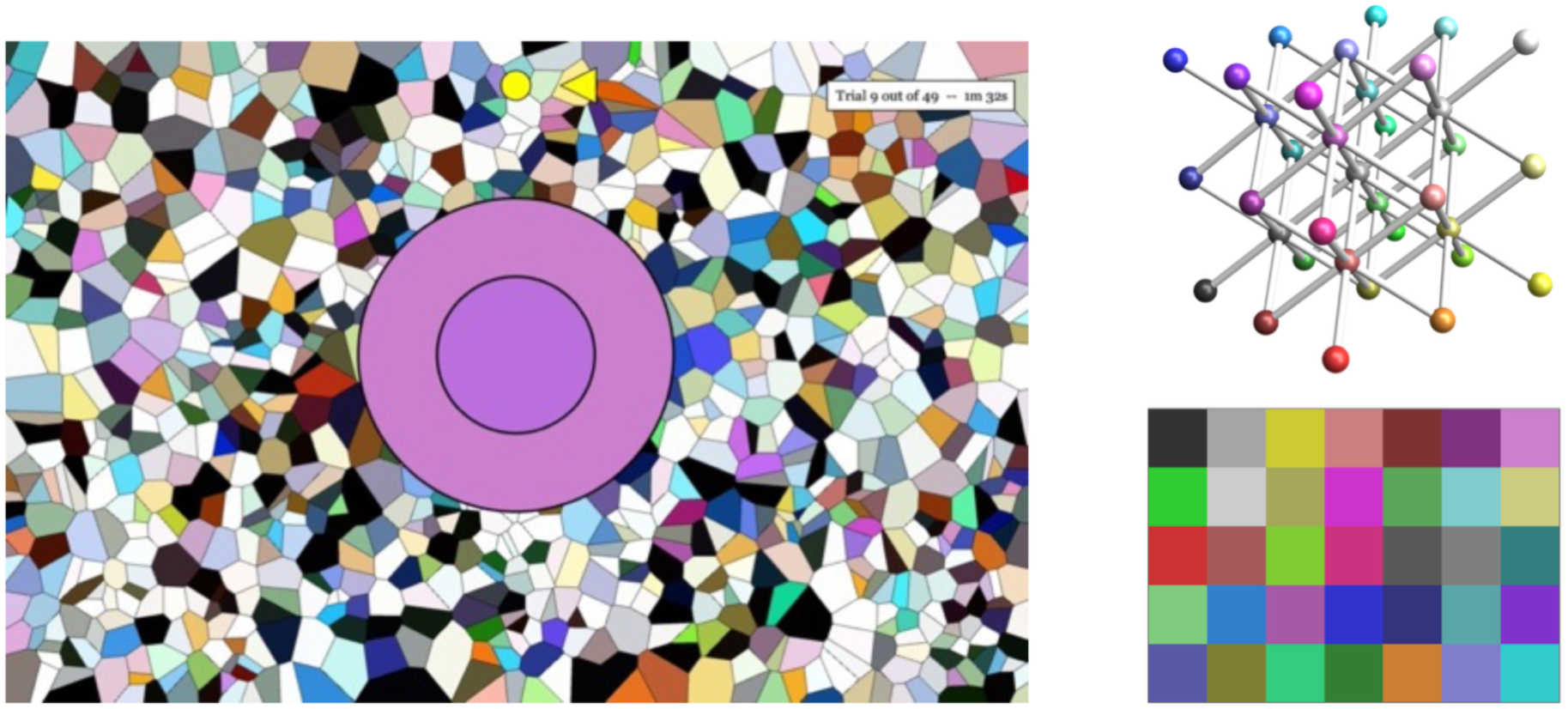
(A) Example stimulus display. A central disk is surrounded by an annulus and presented against a slowly varying colored Voronoi background. The annulus shows the fiducial color, while the central disk is adjusted by the participant. A solid black contour separates disk and annulus to minimize reliance on edge contrast. Directional cues above the stimulus indicate the current adjustment axis. (B) Sampling grid in RGB space. Reference colors were distributed on a body centered cubic lattice within the (0.2, 0.8) interval of the RGB cube. The margin was left to prevent gamut over- und underflows during adjustment. (C) The 35 sampled RGB reference colors used in the experiment.

To estimate the “grain” of RGB space, we sampled 35 reference colors distributed within the RGB cube using a body-centered cubic (BCC) lattice (Figure 2B). The BCC lattice provides high packing efficiency and uniform nearest-neighbor structure, making it well suited for sampling smoothly varying fields in three dimensions. The lattice consisted of eight BCC unit cells confined to the interior of the RGB cube. An empty boundary layer of width 0.2 was left around the cube. This margin provided headroom for overshooting the qualitative criterion and minimized boundary-induced distortions. Overshooting was particularly important for establishing the criterion at the beginning of each trial (see below). The 35 sampled colors are shown in Figure 2C.

### Task

At each reference color, the local grain was measured along seven orientations in RGB space, corresponding to fourteen opposite directions. The seven orientations were chosen to reflect the symmetry of the body centered cubic sampling lattice. They comprise the three Cartesian axes and the four body diagonal directions. The selection provides near isotropic coverage of three dimensional space while remaining experimentally tractable. The seven orientations also constitute a practical compromise. A general ellipsoid in three dimensions has six independent parameters. Fourteen radial measurements provide sufficient constraint for ellipsoidal estimation while keeping the total number of trials feasible.

For each orientation, Notable Qualitative Differences (NQD) were measured in mutually opposite directions. Participants adjusted the comparison color along one direction and then along the opposite direction in a reciprocating manner. They were instructed to set the separation from the reference color such that the difference appeared notably qualitatively different. The specific direction in RGB space was not described to the observers in geometric terms. It was presented simply as one direction and its opposite. NQD regions are approximately an order of magnitude larger than discrimination thresholds and therefore probe perceptual structure at a qualitatively distinct scale. It took observers between 40 and 70 s to finish a single direction, and 5 sessions of 45-60 minutes to finish the whole experiment.

Before recording settings, participants were encouraged to exceed their adopted criterion in both directions at least once. This ensured that the qualitative contrast between the two sides was clearly experienced. For each side, five turning points were recorded. The grain along that orientation was defined as half the difference between the medians of the turning points in the two opposite directions. This procedure exploits the assumed central symmetry of the local grain region and provides an internal consistency check.

Trials were randomized across reference locations and orientations to minimize systematic order effects.

### Construction of Ellipsoidal Regions

For each reference color and each orientation, five turning points were recorded on either side of the reference. The median turning point in each direction was taken as the radial extent of the notably qualitative difference region. The fourteen radial extents define a centrally symmetric set of endpoints in RGB space. Because the local grain region is assumed to be convex and centrally symmetric, we first computed the convex hull of the measured endpoints (Rockafellar, 1970). The minimum volume circumscribed ellipsoid of this hull, also known as the Löwner John ellipsoid (John 1948; Ball 1992; Ben-Tal & Nemirovski, 2001), was then determined. For centrally symmetric convex bodies this ellipsoid is unique. Interior points do not affect the solution, so restricting the computation to the hull preserves the geometry while ensuring numerical stability.

The resulting boundary ellipsoid (Figure 3) is represented by a 3 × 3 symmetric positive definite (SPD) matrix Σ defined through the quadratic form

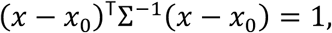

**Figure 3:**
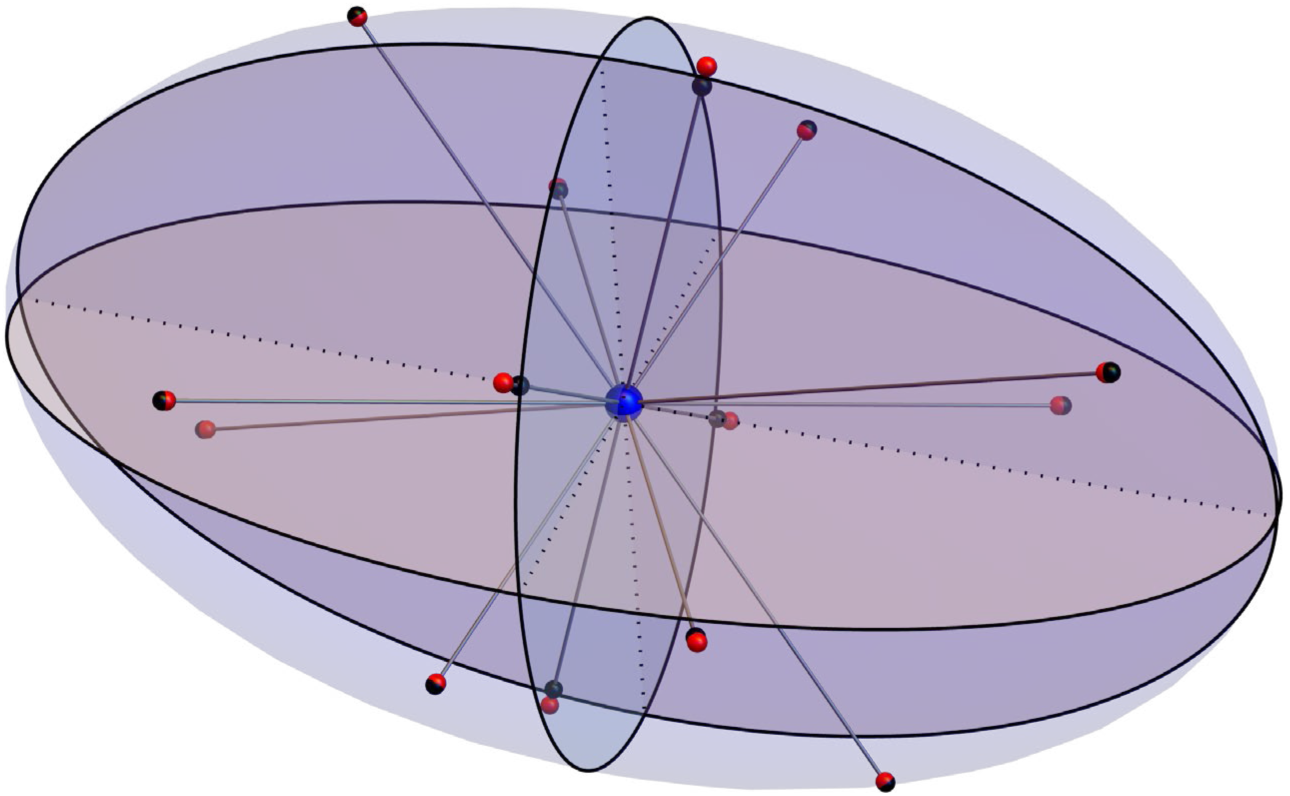
Example of a fitted ellipsoid with principal axes and principal planes. Black spheres indicate the median turning points measured along the sampled directions. Red spheres show the corresponding intersections with the fitted ellipsoid. The close agreement between measured and fitted points indicates that the ellipsoid provides a good representation of the data despite the sparse directional sampling.

where *x*_0_ denotes the reference color. The eigenvectors of Σ determine the principal axes and the eigenvalues correspond to the squared semi axis lengths. Thus each sampled color location is associated with a point in the six dimensional space of symmetric positive definite matrices.

Relative radial deviations between the empirical endpoints and the fitted ellipsoid surface were typically around ten percent and showed no systematic dependence on color location or observer. The settings were also approximately symmetric with a median asymmetry of 10.4%. The ellipsoid therefore provides a compact and stable descriptor of the local grain while replacing small irregularities of the empirical hull by a smoothly structured quadratic form.

### The Metric Field

Once each sampled color is represented by a symmetric positive definite (SPD) matrix Σ, the data define a field in which each location in RGB space is associated with an SPD matrix. To assemble these local measurements into a coherent global field, three operations are required. Ellipsoids must be compared, averaged, and interpolated. All three operations require a metric structure on the space of SPD matrices. Without such a structure, notions of distance, mean, or smooth variation remain undefined. We adopt the Frobenius inner product, which treats symmetric matrices as elements of a high-dimensional Euclidean vector space (see Arsigny et al., 2006; Moakher & Batchelor, 2006; Andreani, Raydan & Tarazag, 2013). For two matrices S and T, the Frobenius inner product is defined as

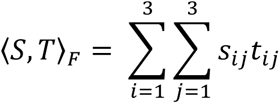

This induces the Frobenius norm

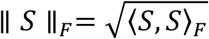

and the corresponding distance

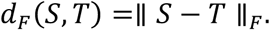

This definition has several decided advantages. It is invariant under rotations of the coordinate frame, so comparisons between ellipsoids are independent of how the RGB space is oriented. It renders the space of symmetric 3 × 3 matrices a six dimensional Euclidean space. Averaging and interpolation therefore reduce to coefficient wise weighted means. This linear structure makes the construction of a global field straightforward and numerically stable. Although alternative Riemannian geometries for symmetric positive definite matrices have been proposed in other fields (Arsigny et al. 2006; Moakher and Batchelor 2006), the present problem does not impose physical constraints. The Frobenius metric therefore provides an economical and transparent choice.

To compare ellipsoids on a common relative scale, we define a dimensionless normalized mismatch measure

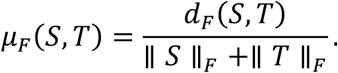

This dimensionless quantity ranges from zero for identical ellipsoids to one for maximal opposition. It captures joint differences in scale, shape, and orientation. The relative Frobenius mismatch was used to assess agreement between different locations within observers, variability across observers, differences between empirical ellipsoids and CIEDE2000-derived ellipsoids, and fidelity of interpolation relative to sampled data.

### Interpolation

The ellipsoids estimated at the 35 sampled locations constitute discrete measurements of local grain. To obtain a metric at arbitrary RGB locations, these samples must be assembled into a continuous field. Because the Frobenius geometry is Euclidean, interpolation reduces to weighted averaging of matrix coefficients.

Let *f* denote an arbitrary point in the RGB cube and let *s_i_* denote neighboring sampled locations. Each sample contributes with a weight determined by its spatial distance from *f*. We construct a partition of unity using Gaussian radial basis functions (Buhmann, 2003):

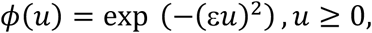

where ε controls the spatial bandwidth.

The weights are defined as

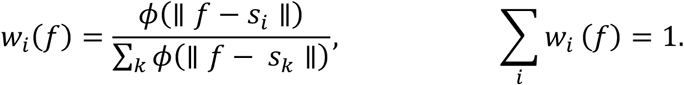

The interpolated ellipsoid at location *f* is then

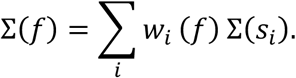

Because the Frobenius geometry is linear, this expression corresponds to a coefficient wise weighted mean of matrix elements.

In practice, interpolation can be restricted to a small neighborhood of sampled points. A six neighborhood corresponding to nearest body centered cubic neighbors is sufficient for most applications. The bandwidth parameter was chosen so that, at a sampled location, the contribution of that location approximately equals the combined contribution of its nearest neighbors. This balances local fidelity and smoothness.

The resulting field varies fairly smoothly across RGB space. At sampled locations, interpolated ellipsoids differ only minimally from empirical estimates. Near the boundary of the RGB cube, interpolation becomes mildly extrapolative due to asymmetric neighborhoods and should therefore be interpreted cautiously, as undue extrapolation might potentially lead to hyperboloids instead of ellipsoids. Overall, the empirical ellipsoids vary fairly smoothly across space. The final metric field reflects the structure of the data rather than artifacts of the interpolation procedure.

## Results

We begin with the empirical ellipsoidal regions at the level of individual observers. Shown at their native scale, the data reveal substantial variability in overall size alongside clear structural regularities. We then quantify agreement across observers and demonstrate that most interindividual differences are multiplicative rather than geometric. After normalizing each observer’s field to a common global scale, the shared spatial structure becomes evident.

On this basis, we construct a group-level representation by computing a Frobenius median ellipsoid at each sampled location. The resulting ensemble defines an overall empirical metric field at the 35 measured points. We characterize its geometric properties, including ellipsoid size, shape distribution, and orientation structure.

We then extend the discrete measurements to a continuous field via interpolation and assess the fidelity of this reconstruction. The resulting field allows dense evaluation throughout RGB space and supports analyses of effective packing, one-dimensional trajectories, planar sections, and other geometric substructures.

Finally, we compare the empirical metric field with the CIEDE2000-derived field, highlighting both substantial agreement in global structure and systematic differences in local anisotropy.

### Initial Structure and Observer Variability

At first inspection, the raw ellipsoidal regions differ markedly across observers. The most conspicuous variation concerns overall size. Some observers adopt a conservative criterion, yielding small grain regions, whereas others adopt a more liberal criterion. When ellipsoids are plotted at true scale, this spread is substantial. For reference, the resulting grain radii corresponded roughly to 5 ± 2 CIEDE2000 units. Despite these scale differences, the spatial pattern across RGB space appears similar. Locations that exhibit large grain for one observer tend to do so for others as well. Visual inspection therefore suggests that interobserver variability is largely a scaling rather than structural. Figure 4 shows the raw ellipsoids for all eight observers at true scale. The range of overall sizes is striking, yet similarities in spatial variation are evident even before normalization.

**Figure 4:**
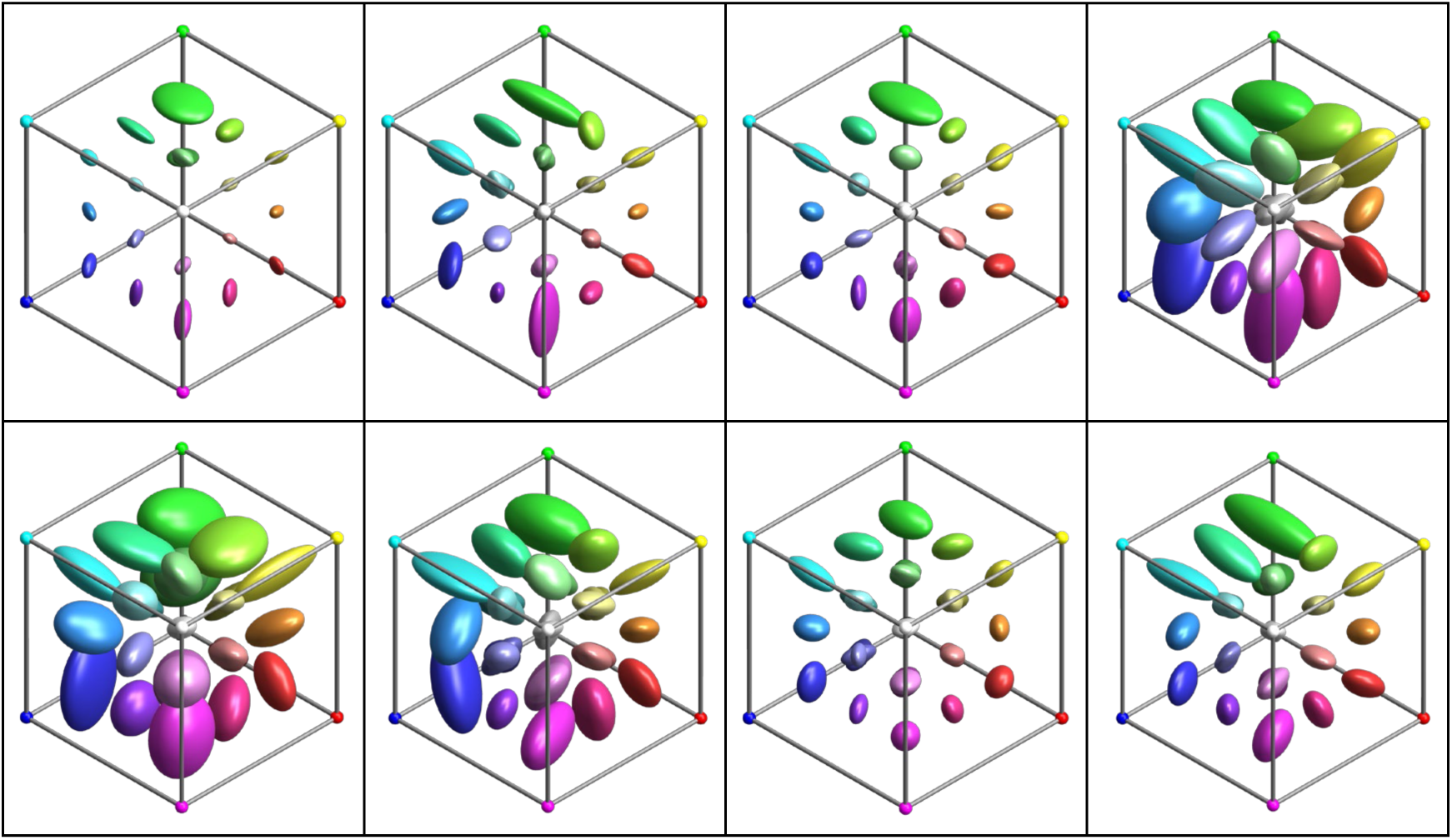
The raw data (one particular view) of the 8 participants. Ellipsoids drawn at true size. Not all 35 ellipsoids are visible due to occlusion.

To quantify this observation, we computed correlations between ellipsoid sizes across observers. Different measures of ellipsoid size (trace, volume, diameter, Frobenius norm, linear size) all correlated highly and significantly. Figure 5 shows the Spearman correlation matrix for the cube root of ellipsoidal volume. The median correlation was 0.82, with a range from 0.70 to 0.86 (all p < 0.001). Thus observers agree reliably on where grain is relatively large or small, even though they differ in absolute scale. There was no difference between the four (old) authors and the four (young) naïve observers. The median correlation within groups (0.80) was essentially the same as that of the between group correlations (0.82).

**Figure 5:**
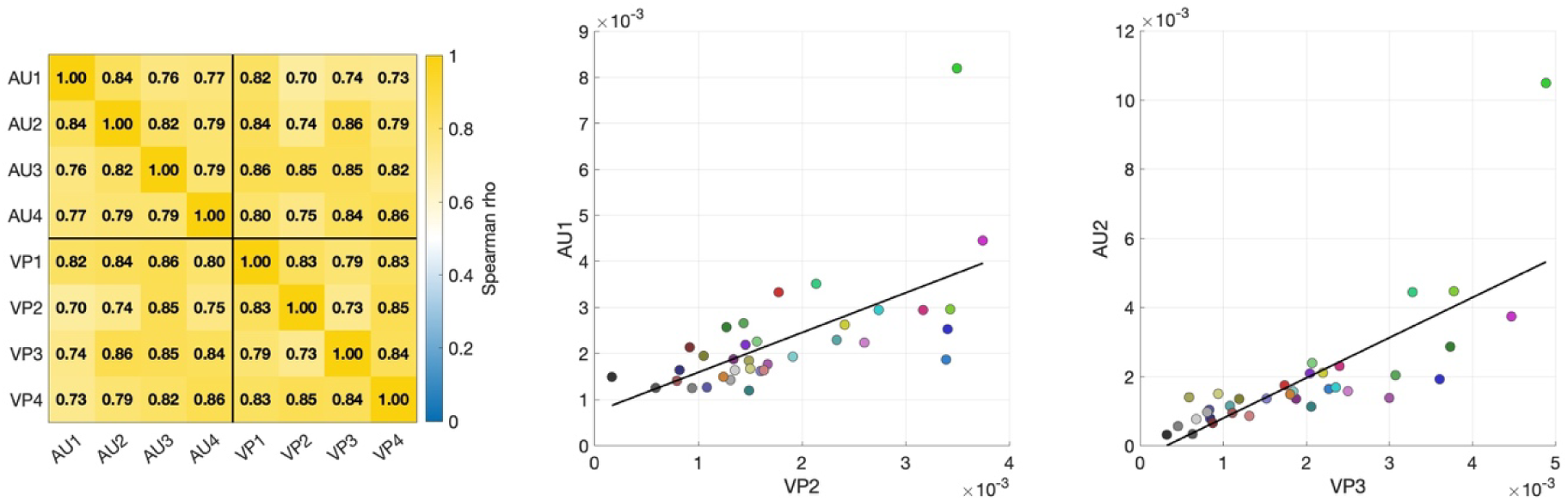
Inter-observer correlation matrix for the cube root of ellipsoid volume, together with the pairs showing the lowest (0.70) and highest (0.86) correlations.

Across observers, median ellipsoid diameters varied by approximately a factor of four. Given the high rank agreement, this variation is most parsimoniously interpreted as reflecting individual criterion levels rather than differences in the underlying spatial structure of the metric field.

### Normalization

To isolate the shared spatial structure, we normalized each observer’s ellipsoids by scaling their median Frobenius norm to a common reference value in line with the results of the observer with the most conservative criterion who produced the smallest ellipsoids. This procedure removes global criterion differences while preserving the relative pattern of variation across RGB space within each observer. After normalization, the fields become markedly more homogeneous. Differences in absolute scale largely disappear, revealing a common spatial organization that was already latent in the raw data. Figure 6 shows the normalized ellipsoids for all observers. The improvement in comparability is immediately apparent. Locations that exhibited similar relative grain before normalization now align closely in absolute size as well. Residual differences remain, but they are modest compared to the original spread.

**Figure 6:**
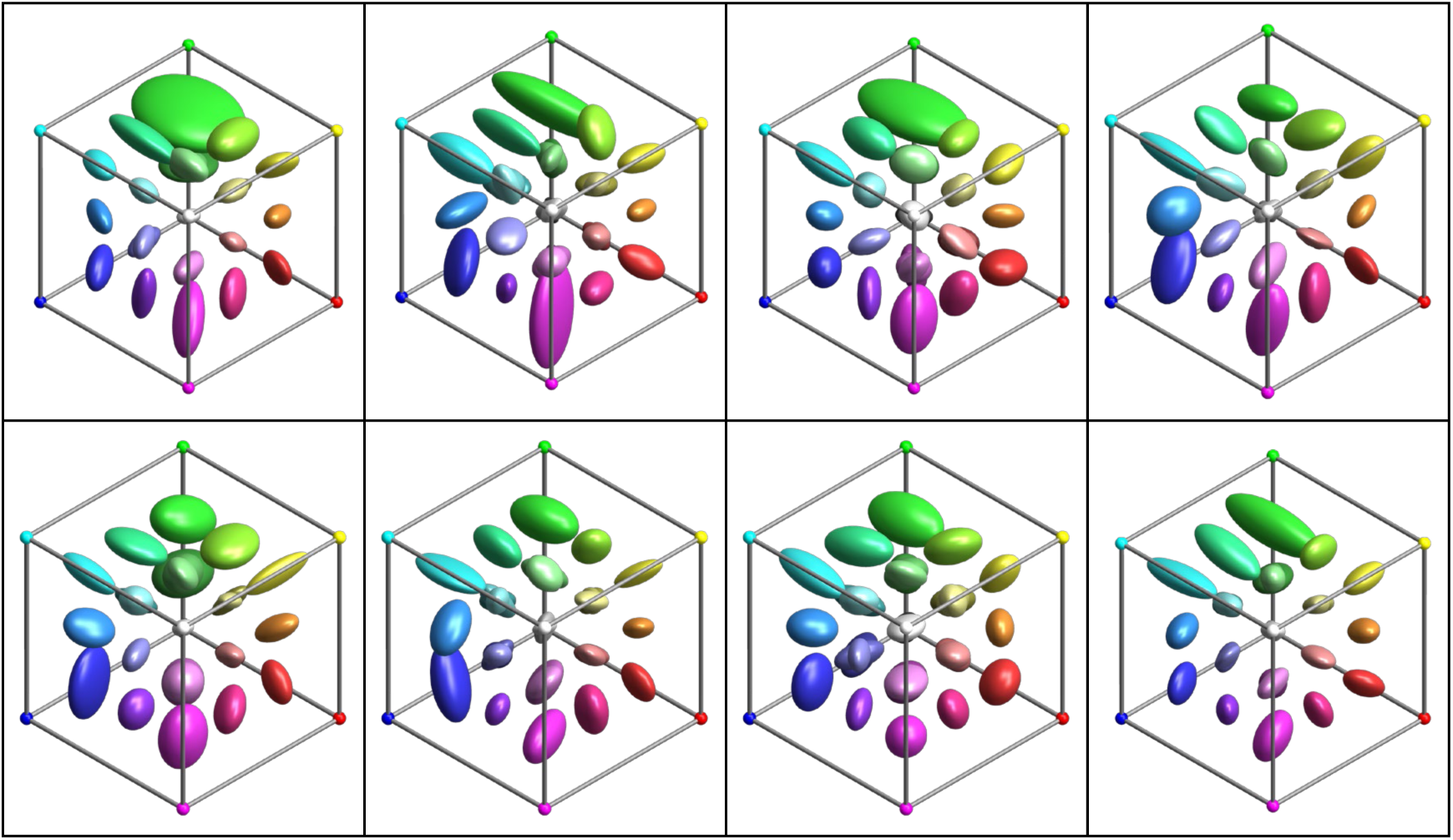
Normalized grain fields for all eight observers. Ellipsoids are shown at double size for visibility. Each observer’s field has been scaled so that the median Frobenius norm matches a common reference value. Compared with the raw data in Figure 4, differences in global size are largely eliminated. The remaining variation reflects genuine structural differences rather than criterion level. Spatial gradients and systematic anisotropies are more readily apparent after normalization. Note that not all 35 ellispoids are visible to to occlusion.

Spatial gradients across the RGB cube become more clearly visible after normalization. In particular, systematic increases along the achromatic direction and chromatic asymmetries can now be seen without the confound of global scale differences.

Importantly, normalization does not alter rank-order relations across locations. The Spearman correlations reported above remain unchanged, since scaling by a positive constant preserves ordering. Thus the procedure removes only a global scale factor while leaving the geometric structure intact.

To give a visual sense of what the Frobenius mismatch means geometrically, Figure 7 displays ten real observer–location cases arranged in order of increasing mismatch (value printed in each panel). In every panel the grey wireframe is the median field’s ellipsoid at that location and the translucent coloured surface is the individual observer’s ellipsoid, drawn in the location’s own RGB color; all ten panels share a common scale, and the per-observer global scale has been removed. The sequence spans most of the observed range, from close agreement (≈0.07) through the typical between-observer level (≈0.20) to the largest departures (≈0.38).

**Figure 7.**
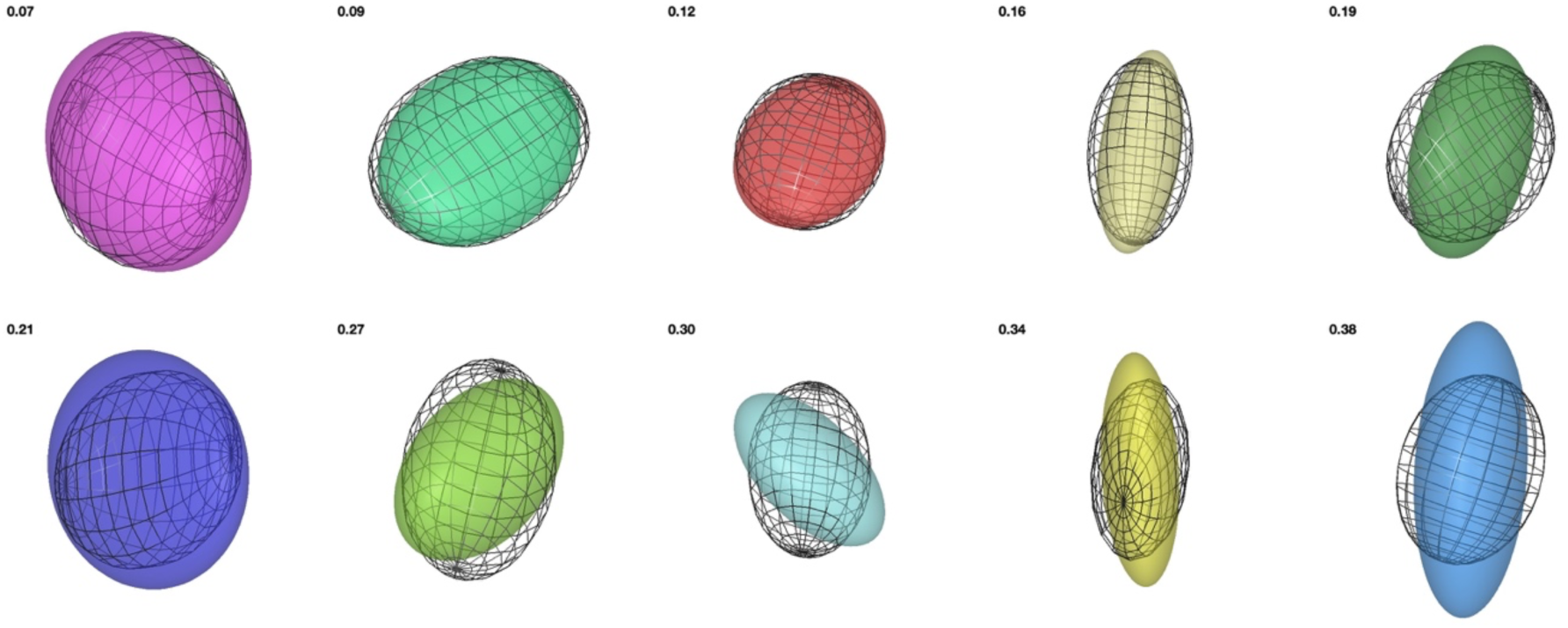
Visual calibration of the Frobenius mismatch using ten real observer–location cases, arranged in order of increasing mismatch (2 × 5 panels, value printed at the top-left of each). In every panel the grey wireframe shows the median field’s ellipsoid at that location and the translucent coloured surface shows the individual observer’s ellipsoid, drawn in the location’s RGB colour; all panels share a common scale, so relative sizes are preserved and the per-observer global scale has been removed. The sequence spans the bulk of the observed range, from close agreement (≈0.07) through the median between-observer level to large departures (≈0.38), giving a direct visual reference for the mismatch values reported elsewhere. The apparent agreement is view-dependent: an ellipsoid can often be rotated to coincide closely with the median, so these fixed orthographic views (and the Frobenius mismatch itself) capture orientation differences that a freely-rotated comparison would hide.

Even at the upper end of this range the two ellipsoids remain similar in size and overall shape, differing mainly in the length of one axis or a modest tilt of the principal directions. This supports the interpretation that the qualitative grain structure is shared across observers up to a global scale factor, with interobserver variability that is primarily criterion-based rather than structural.

### The Overall Empirical Field

Given the high degree of homogeneity after normalization, it becomes meaningful to aggregate across observers. At each sampled RGB location, we computed a Frobenius median of the normalized ellipsoids. Concretely, this corresponds to taking the coefficient-wise median of the symmetric positive-definite matrices. The resulting set of ellipsoids constitutes the overall empirical field at the 35 sampled locations. Figure 8 shows this group-level metric field from three orthogonal viewing directions.^1^ The spatial variation of ellipsoid size and orientation is smooth across the RGB cube. No abrupt discontinuities are visible, and local changes occur gradually from one location to the next. The existence of a coherent group field therefore justifies the subsequent construction of a continuous metric field through interpolation. The complete set of parameters for the 35 ellipsoids is given in Appendix A2.

**Figure 8.**
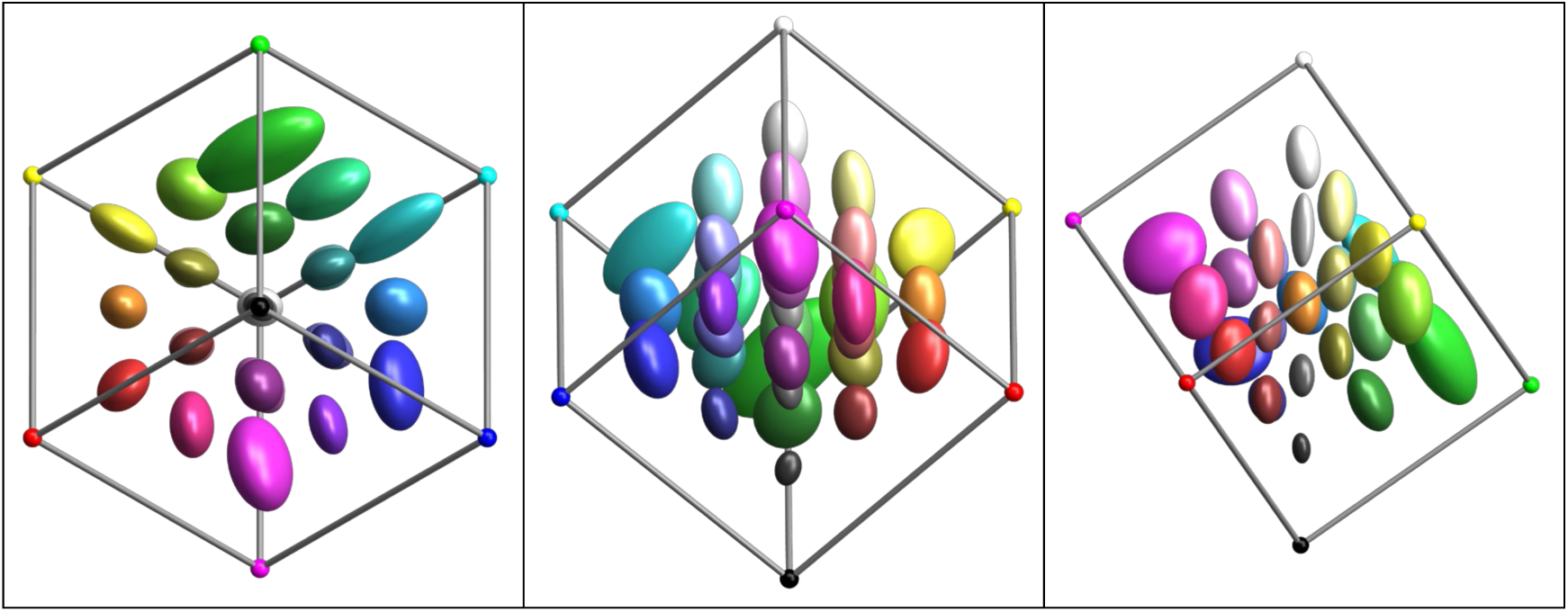
Group-level metric field obtained as the Frobenius median of the normalized individual fields shown in Figure 6. Ellipsoids are displayed at double size for visibility. Three orthogonal views are provided. Left panel is viewed along the white–black axis, center panel along the purple–green axis, and right panel along the orange–teal axis. The field varies smoothly in both size and orientation across RGB space.

To quantify the agreement between the individual normalized fields and the group representation, we computed relative Frobenius mismatches between each observer’s ellipsoids and the corresponding median ellipsoids. The quartiles were {0.16, 0.21, 0.27}, indicating tight clustering around the group structure. Figure 9 decomposes this overall variability (leftmost panel) into differences of volume, elongation and orientation. In each panel the solid orange line marks the typical difference between locations, that is, the geometric variability across RGB space for the same statistic.

**Figure 9:**
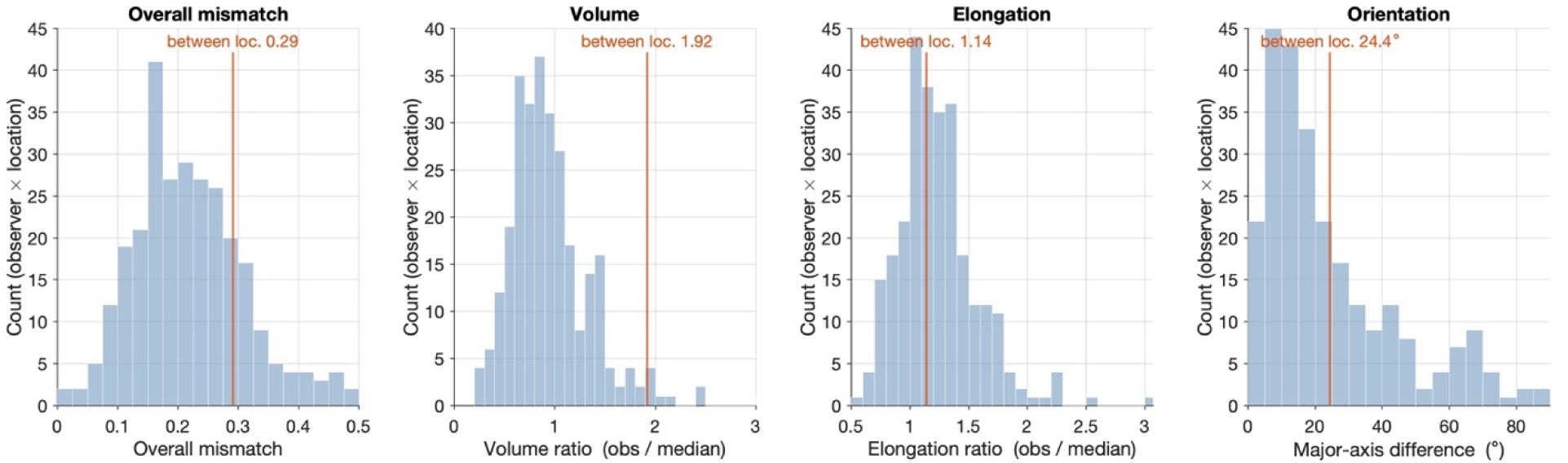
Geometric differences between each observer’s threshold ellipsoid and the median field at the same location, pooled over all observers and locations (8 × 35 = 280 pairs, representation with per-observer global scale removed). Left: overall normalised Frobenius mismatch between the observer and median ellipsoids (median 0.21). Centre-left: ratio of ellipsoid volumes, √(det A / det B) (median 0.89). Centre-right: ratio of elongations √(λ₁/λ₃) (median 1.19). Right: 3-D angle between the major axes, excluding near-spherical cells with ill-defined major axis (λ₁/λ₂ < 1.25; 254 of 280 pairs; median 17.1°). Solid orange lines mark the corresponding median difference between locations (0.29, 1.92, 1.14, 24.4°), the typical magnitude of the stimulus effect for each statistic.

Across observers and locations the ellipsoids cluster around the median in both local size and elongation (volume ratio median 0.89, elongation ratio median 1.19). For orientation most ellipsoids are nearly co-oriented with the median (major-axis difference median 17.1°), with a tail of larger deviations. Overall, the median field provides an accurate summary of individual observer structure. The interobserver differences are small relative to the variability across RGB space for overall shape (0.21 vs 0.29) and, most strikingly, for volume: the median ellipsoids differ in size across locations by roughly a factor of two (between-location ratio 1.92), far exceeding the ∼10% volume deviation of individual observers from the median. Orientation shows the same ordering (between-location 24.4° vs 17.1°), with a few large major-axis differences, as expected when the two largest axes are similar in length and the principal direction is correspondingly ill-determined. For elongation the interobserver spread (1.19) is comparable to, and slightly exceeds, the between-location variation (1.14), reflecting that elongation changes little across the gamut.

The geometry of the group-level ellipsoids is not arbitrary. Both shape and orientation exhibit systematic structure across the 35 sampled locations. Figure 10 summarizes these statistics for the median observer. Each threshold ellipsoid is characterized by the eigenvectors and eigenvalues λ₁ ≥ λ₂ ≥ λ₃ of its covariance matrix, where the eigenvalues are the squared lengths of the principal semi-axes.

**Figure 10.**
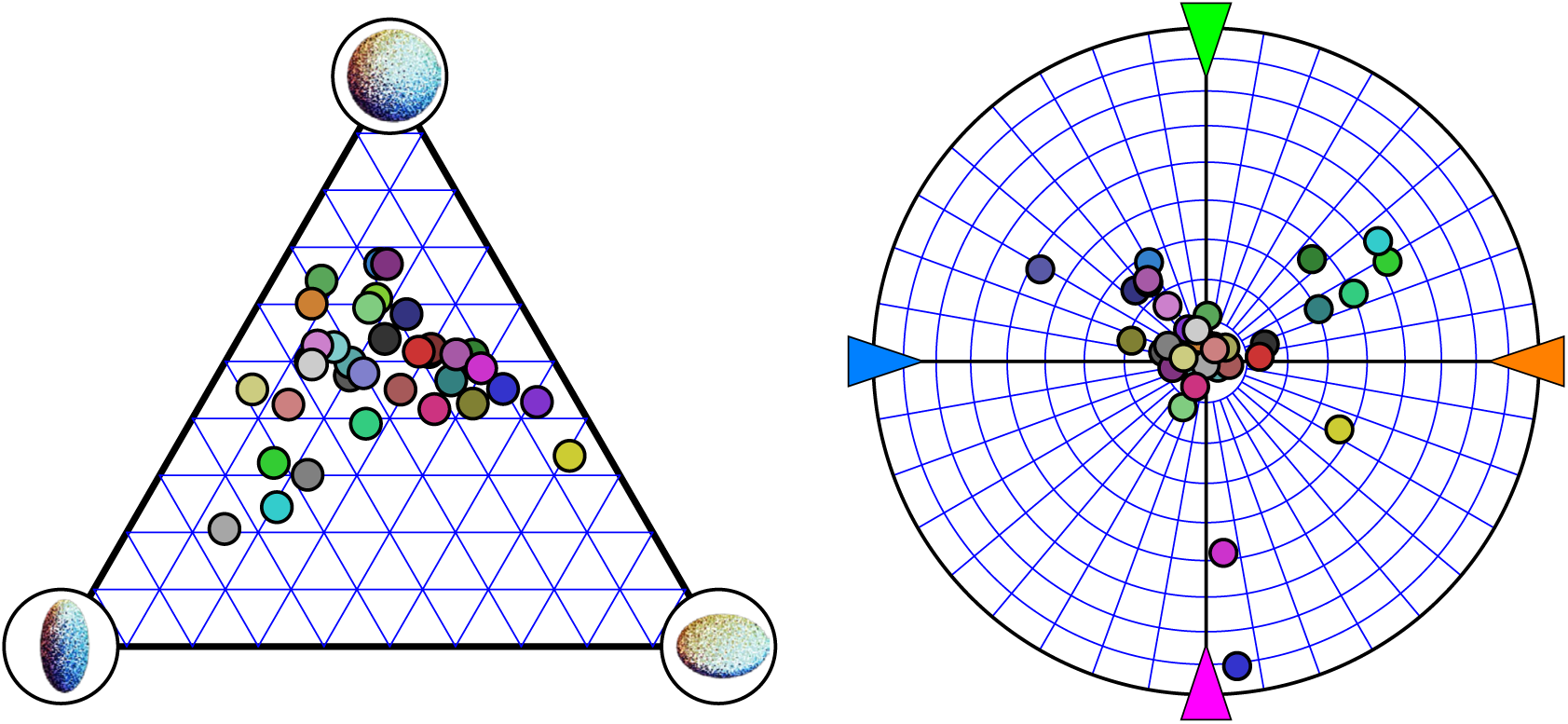
Shape and orientation statistics of the group-level ellipsoids across the 35 sampled locations. Left panel shows the distribution of ellipsoid shapes within a triangular representation spanning spheroid, prolate, and oblate forms. Most ellipsoids are near spheroidal with a slight bias toward prolate shapes. Right panel shows the orientation of the major axes in an area-correct polar projection with the achromatic direction at the center. Only one hemisphere is shown. The radii indicate 10 deg steps in elevation. Orientations cluster strongly near the achromatic axis, with smaller systematic deviations toward chromatic directions.

The left panel shows the distribution of ellipsoid shapes in a triangular representation (Westin ternary plot) spanning spheroidal, prolate, and oblate forms. Shape is captured by three coordinates that are normalized by the eigenvalue sum λ₁+λ₂+λ₃, so that they add to one and are independent of overall size: a prolate index (λ₁−λ₂)/(λ₁+λ₂+λ₃), an oblate index 2(λ₂−λ₃)/(λ₁+λ₂+λ₃), and a spherical index 3λ₃/(λ₁+λ₂+λ₃). The three corners of the triangle correspond to the limiting forms: prolate (cigar-shaped, one long axis), oblate (disc-shaped, one short axis), and spherical (isotropic). Most ellipsoids lie close to the spheroidal region, with a modest but consistent bias toward prolate shapes.

The right panel shows the orientation of the major axes (the first eigenvector of each ellipsoid) in an area-correct (Lambert equal-area) polar projection, with the achromatic direction [1,1,1] at the center. The radial coordinate is the angle of the major axis away from the achromatic axis, so that the center corresponds to a major axis aligned with the achromatic direction and the bounding circle to a major axis lying in the chromatic plane; the angular coordinate gives the chromatic azimuth, with the principal chromatic directions (orange, blue, green, and magenta) marked around the rim. Orientations are displayed for one hemisphere only, since antipodal directions are equivalent. The majority of major axes cluster near the achromatic axis. Smaller but systematic deviations are visible toward secondary chromatic directions.

This pattern is highly structured and far from the isotropic distribution expected from a rotationally invariant random SPD ensemble such as the default Wishart distribution (Wishart, 1928).

The grain field is therefore structured not only in overall scale but also in anisotropy and orientation. The geometry of local qualitative equivalence regions reflects systematic organization within RGB space rather than random variability. This structured organization suggests that a smooth global metric field can be reconstructed from the sparse local measurements.

### Construction of the Continuous Metric Field

At this stage several conclusions follow directly from the empirical measurements. First, the ellipsoidal approximation is adequate. Relative deviations from the convex hull are small and show no systematic dependence on location. Second, interobserver differences are largely multiplicative. After normalization, spatial structure aligns closely across observers. Third, the aggregated ellipsoids vary smoothly across RGB space. No abrupt discontinuities are visible in the sampled locations. Fourth, anisotropy is systematic. Ellipsoid orientations cluster along a small number of recurring directions rather than being randomly distributed. Taken together, these findings justify treating the 35 empirical ellipsoids as samples from an underlying smooth metric field in RGB space.

The normalized and aggregated ellipsoids at the sampled locations form the empirical backbone of the metric field. Using the interpolation procedure described in Methods, these discrete samples can be extended to arbitrary locations within the RGB cube. The result is a smooth field of symmetric positive-definite matrices defined over RGB space. At every location, the field assigns an ellipsoid representing the local region of notable qualitative equivalence.

This step marks a conceptual transition. The experiment provides measurements at 35 discrete locations. Interpolation converts these measurements into a continuous geometric object. Once constructed, the field permits evaluation of local grain at any location in RGB space without performing additional experiments. In this sense, the metric field bridges empirical measurement and theoretical representation.

To assess the fidelity of interpolation, we compared the interpolated ellipsoids at the sampled locations with the original empirical ellipsoids in Figure 11.

**Figure 11.**
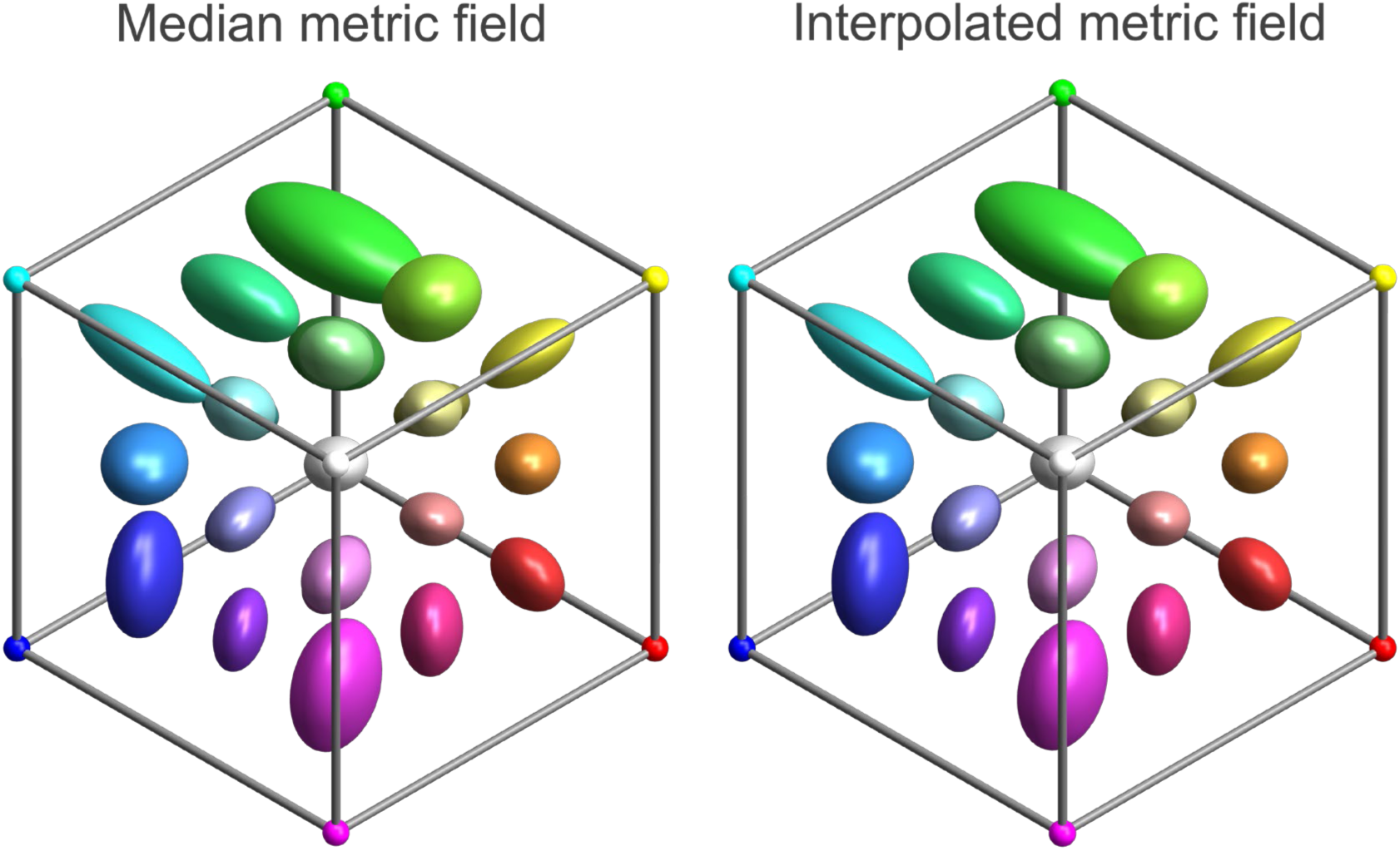
At left the median grain at the basis locations from Figure 8. At right the interpolated grain at the same locations. With careful inspection, a minor discrepancy in ellipsoid orientation can be detected at the saturated green corner of the cube. The interpolation algorithm essentially leaves the grain at the basis locations intact. Note that the interpolation assigns a weight of 0.5 to the measured ellipsoid at the target location, so the interpolated estimate is not entirely independent of the measurement, although differences in orientation and axis ratios are in principle possible.

The relative Frobenius mismatches between empirical and interpolated ellipsoids were extremely small (quartiles {0.017, 0.025, 0.035}). Even the worst case remained well within the variability observed across observers. Visually, differences are difficult to detect. This indicates that interpolation introduces only minimal distortion. The smooth field is not imposed upon noisy data; rather, the empirical data themselves are already slowly varying.

A stricter leave-one-out test, in which each location is removed and then reconstructed from the remaining 34, naturally yields larger mismatches (quartiles {0.11, 0.14, 0.20}), since the target ellipsoid no longer contributes to its own estimate. Even so, the median leave-one-out mismatch of 0.14 stays well below the typical between-observer variability of about 0.21, confirming that any single location is well predicted by its neighbors. The largest errors occur at the edges and corners of the sampled gamut, where a held-out location can only be extrapolated from one-sided neighbors.

Near the boundary of the RGB cube, interpolation necessarily merges into extrapolation, because neighboring sample points are confined to one side. In these regions, effective support shrinks to a small subset of interior samples. As a result, extrapolated grain values tend to stabilize. The boundary layer (approximately 0.2 in thickness) occupies a non-negligible fraction of volume. However, its thickness is on the order of the grain size itself. For most practical applications involving interior colors, boundary effects are therefore unlikely to dominate. A more detailed characterization of boundary regions would require dedicated measurements or the development of extrapolation procedures tailored to edge geometry, both of which are beyond the scope of the present study.

Several conclusions follow from the construction and validation of the continuous metric field. First, the empirical grain structure is intrinsically smooth. The aggregated ellipsoids vary gradually across sampled locations, and interpolation introduces only negligible deviation. Second, anisotropy is systematic rather than random. Ellipsoid shapes and orientations cluster along a small number of recurring directions and vary smoothly across RGB space. Third, interobserver variability is primarily multiplicative. After normalization, the geometric structure of the field is highly consistent across participants. Fourth, sparse sampling suffices to recover the large-scale organization of the metric. The 35 measured locations provide enough information to reconstruct a coherent global field over RGB space.

Interpolation reproduces the empirical ellipsoids at sampled locations with minor difference. This allows the field to be evaluated densely throughout the interior of the RGB cube. The experiment provides a finite set of anchor points from which a volumetric perceptual geometry can be constructed. We can make virtual measurement of local grain at any location of theoretical interest, without additional data collection.

A direct consequence of this construction concerns the effective granularity of RGB space. If each ellipsoid represents a region of qualitative equivalence, then the continuous field defines a tiling of the cube by overlapping grain regions. Dense evaluation makes it possible to estimate how many such regions the cube can accommodate, an issue to which we return below. In effect, the discrete ellipsoids are sufficient to define a continuous perceptual metric field. This field captures both scale and anisotropy of qualitative color differences and can be evaluated at arbitrary locations within the RGB cube.

To visualize the interpolated metric field as a volumetric object, we evaluated it on a dense body-centered cubic lattice within the RGB cube. This procedure does not introduce new empirical data. It simply queries the continuous field at a large number of interior locations. The resulting ensemble of ellipsoids reveals a coherent three-dimensional geometry. Ellipsoid size and orientation vary smoothly throughout the cube, without abrupt distortions or artifacts. Regions of gradual expansion and contraction are clearly visible, and anisotropies evolve continuously across space.

### One Thousand Colors in RGB

Because the field is defined everywhere in the interior, one may conceptually regard the RGB cube as filled with overlapping regions of qualitative equivalence. This makes it possible to address a natural question: how many such regions can the cube accommodate? Figure 12 illustrates the idea. The cube is filled with local grain ellipsoids placed on a body-centered cubic lattice whose spacing is set to the median grain size, each ellipsoid interpolated from the metric field and colored by its location. At this spacing the grains approximately tile the space, some leaving small gaps and others overlapping, so the arrangement sits close to a practical packing limit. Across the full cube the lattice contains 1241 grains.

**Figure 12.**
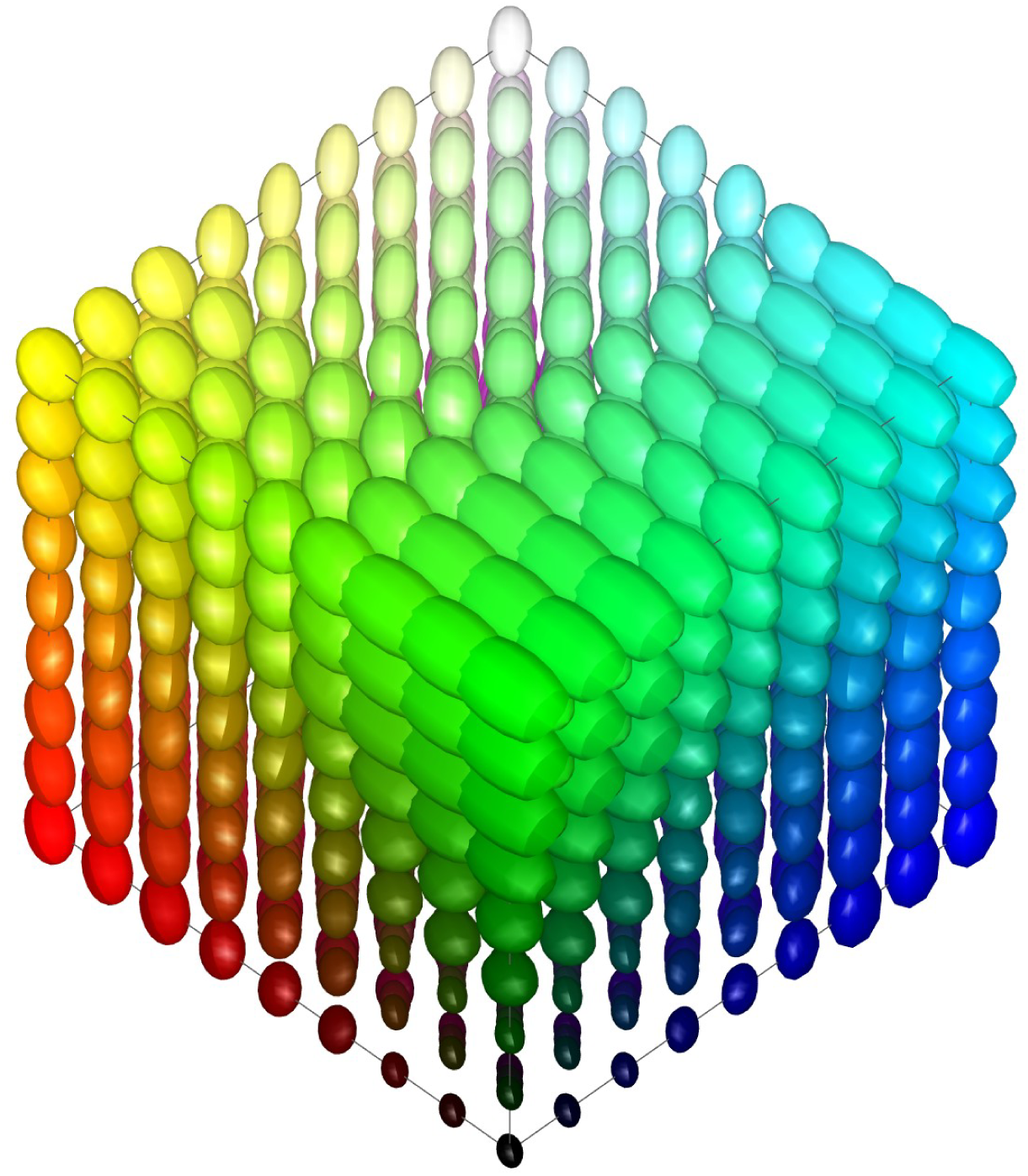
The RGB cube filled with local grain ellipsoids on a body-centered cubic lattice whose spacing equals the median grain size, each ellipsoid colored by its location and interpolated from the median metric field. At this density the grains approximately tile the cube, with some overlap and small gaps, illustrating the approximate number of qualitatively distinct regions the cube can accommodate.

We can make this estimate quantitative by integrating the metric field itself. Treating each grain as a cell whose volume is set by the local metric and applying a realistic packing fraction, the unit cube accommodates on the order of one to two thousand grain regions, consistent with the body-centered cubic arrangement of Figure 12. This should be read as an order of magnitude rather than an exact count, since grain size varies across space and the ellipsoids are neither identical nor regularly arranged. The estimate also depends strongly on viewing conditions. State of adaptation, spatial separation of the stimuli, surround context, and other aspects of the experimental arrangement can substantially alter discrimination performance (for review, see Mollon & Danilova, 2026). Any count of discernible colors should therefore be regarded as conditional on a particular viewing configuration rather than as a universal property of human vision.

A figure of one thousand is strikingly smaller than the classical estimates of the number of discriminable colors, which often fall in the million range (Judd & Kelly, 1939; Nickerson & Newhall, 1943). Pointer and Attridge (1998) obtained 2.28 million by counting unit CIELAB cubes within the optimal-color solid under D65. These estimates are not contradictory. Because color space is three-dimensional, the number of distinguishable regions grows as the cube of the reciprocal of the counting unit, so adopting a finer unit inflates the total steeply: the millions obtained from just-noticeable differences and the roughly one thousand qualitative grains are the same space measured at different resolutions, with a single grain spanning on the order of ten threshold steps along each axis.

### Global Exploration of the Metric Field

Having constructed a continuous perceptual metric field, we now examine its structure along geometrically interesting submanifolds of RGB space. Although the analyses are presented in the original sRGB measurement space, the corresponding metric field can be transformed directly into linear RGB, XYZ, LMS, DKL, and other color-space representations (Appendix A3), allowing the same geometric structure to be examined in alternative coordinate systems. These analyses do not introduce new data. They evaluate the interpolated field at locations of particular perceptual relevance. We begin with the achromatic axis, then consider opponent chromatic directions, and subsequently turn to circular and planar sections.

The achromatic axis connects black and white within the RGB cube. It represents variation in luminance without change in chromatic balance and therefore provides a fundamental reference direction. Figure 13 shows interpolated ellipsoids sampled along the grayscale axis. The major axes of the ellipsoids tend to align with the achromatic direction. This alignment indicates greater tolerance for variation along the luminance dimension than for chromatic deviation in neutral colors. As noted by Sharpe and Wyszecki (1976), stimulus separation and surround configuration can preferentially elevate achromatic thresholds, potentially affecting the absolute grain sizes reported here. Grain volume increases monotonically from black toward white.

**Figure 13.**
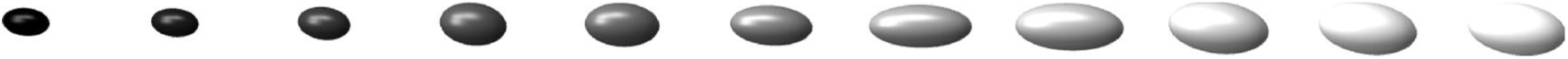
Interpolated ellipsoids sampled along the grayscale axis. The major axes tend to align with the achromatic direction. Grain volume increases monotonically from black to white.

The increase is gradual and continuous. No reversals or local extrema are observed within the sampled interior. Thus, within the present framework of qualitative grain, lighter neutral colors occupy larger regions of equivalence than darker ones.

We next examine the four body diagonals of the RGB cube. In addition to the achromatic axis, these include three chromatic diagonals corresponding to red–cyan, yellow–blue, and purple–green opponent directions. Figure 14 plots grain volume along these diagonals, expressed relative to the median volume of the field. Along the achromatic axis, the monotonic increase described above is confirmed quantitatively.

**Figure 14.**
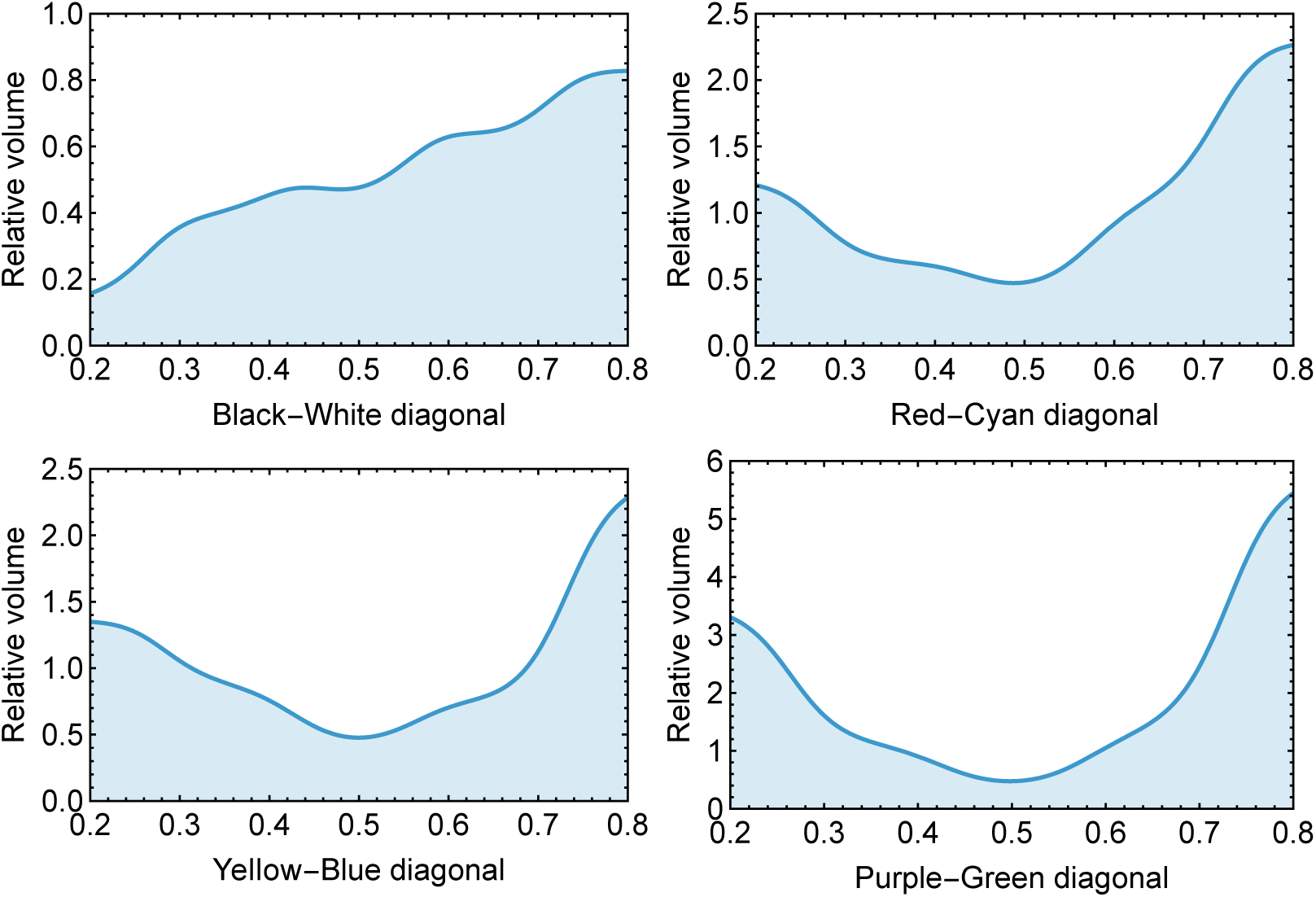
Relative grain volume along the four body diagonals of the RGB cube. Along the achromatic axis, grain volume increases monotonically from black to white. Along the chromatic diagonals, grain volume exhibits V-shaped profiles with minima near the cube center and asymmetric increases toward the boundaries. Near the boundary layer, extrapolated values stabilize due to reduced interpolation support.

**Figure 15.**
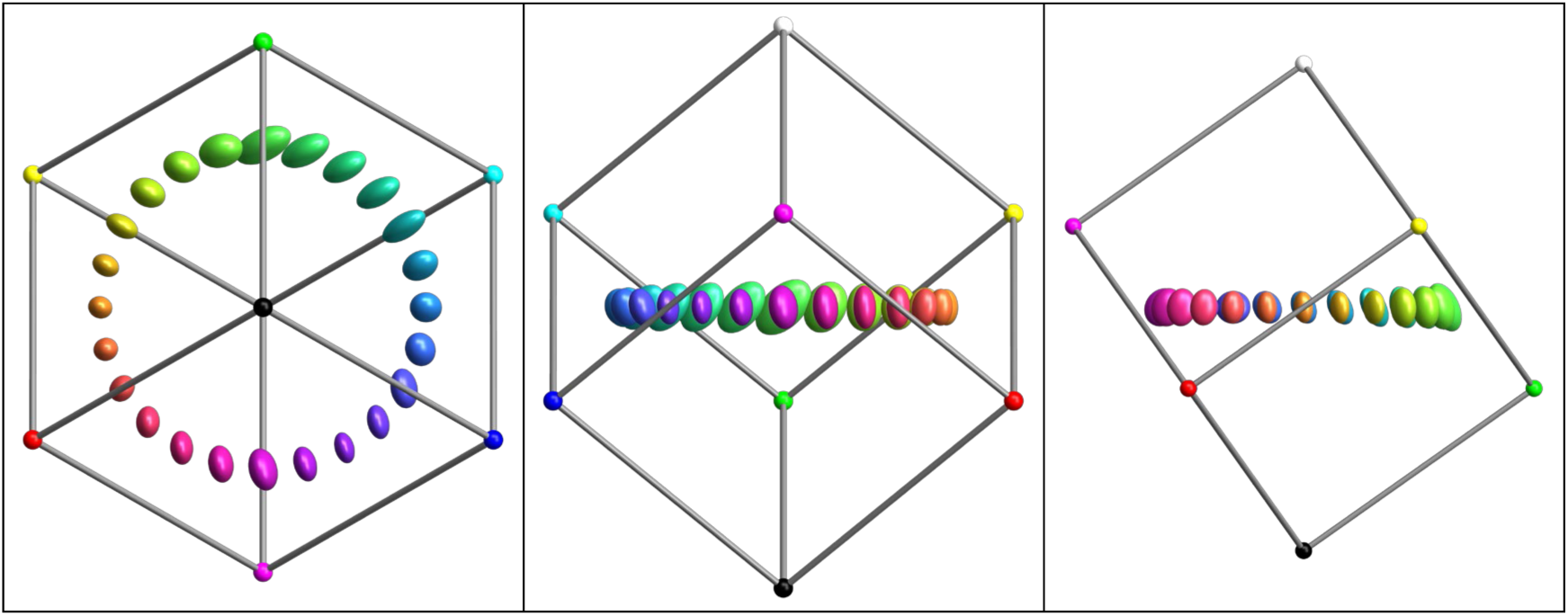
Interpolated ellipsoids sampled along a circular trajectory in RGB space. Ellipsoids are shown at true size. A true circle in RGB space was chosen to avoid extrapolation and boundary-induced distortions associated with the locus of most vivid colors.

Along the chromatic diagonals, a different pattern emerges. In each case, grain volume exhibits a minimum near the center of the cube and increases toward both ends of the diagonal, producing a V-shaped profile. These functions resemble more conventional color discrimination functions measured along opponent axes, such as those reported by Krauskopf and Gegenfurtner (1992), although the present metric concerns finite qualitative differences rather than just noticeable differences.

These V-shaped profiles are not symmetric. Along the red-cyan diagonal the volume increases more steeply at the cyan side. Along the yellow-blue diagonal the volume increases more steeply at the blue side. Along the purple-green diagonal the volume increases more steeply at the green side. Thus there exists an unbalance between the blue–cyan-green and the purple-red-yellow parts of the RGB volume. Similar asymmetries have been discussed in the context of warm and cool colors (Albertazzi, Koenderink, & van Doorn, 2015; Specker et al., 2020; Knoblauch, Werner, & Webster, 2023; Manalansan, Whitehead, & Webster, 2025; Koenderink, van Doorn, & Braun, 2024a). While the present data do not identify the underlying cause, a possible explanation is ecological rather than physiological. Natural illuminants and surface reflectances are strongly constrained and are often well described by a small number of parameters, particularly overall intensity and spectral slope (Koenderink, 2010).

Variations in spectral slope correspond closely to the familiar warm–cool dimension. If perceptual coding is adapted to these dominant regularities of the visual environment, asymmetries in discrimination geometry might naturally emerge along corresponding directions in color space. Whether the grain asymmetry reported here reflects such environmental statistics, specific neural mechanisms, or a combination of both remains an open question.

To examine variation across hue within the interior of RGB space, we evaluated the metric field along a closed circular trajectory. Rather than following the locus of most vivid colors at the boundary of the cube, which forms a twisted polygonal loop, we chose a true circle embedded within the interior of RGB space. This avoids extrapolation and ensures a smooth progression that is not influenced by the non-smooth geometry of the cube boundary.

Figure 15 shows interpolated ellipsoids sampled along this circle. Because the path lies entirely within the interior, the ellipsoids reflect the intrinsic structure of the metric field rather than boundary artifacts. Size and orientation vary continuously along the trajectory.

To quantify variation along the circle, we computed grain volume as a function of position along the path. Figure 16 shows the resulting profile. Grain size varies markedly around the circle. The portion of the path corresponding to hues that admit dominant wavelengths, including the blue region, exhibits a structured modulation that is qualitatively reminiscent of classical wavelength discrimination curves. A similar non-uniformity is evident in modern hue discrimination measurements, such as those reported by Witzel and Gegenfurtner (2015, see their Figure 3). Although those studies concern threshold-level discrimination, whereas the present measure concerns finite qualitative steps, both approaches reveal systematic variation across hue.

**Figure 16.**
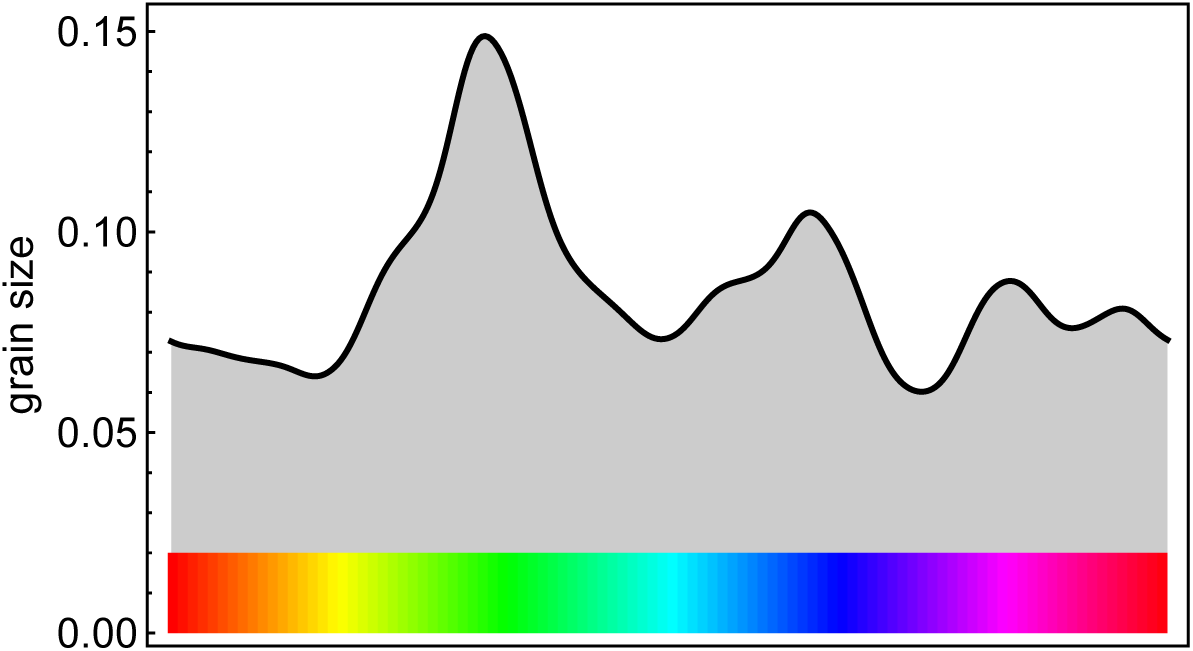
Grain volume along the interior chromatic circle shown in Figure 15. The left portion of the path corresponds to colors admitting dominant wavelengths and may be compared qualitatively with classical wavelength discrimination functions. Grain varies substantially around the circle, confirming that the chromatic manifold is not uniform.

The chromatic manifold is therefore clearly not uniform. Equal angular steps in RGB space do not correspond to equal qualitative steps. In this sense, the circle is not “well tempered,” consistent with the phenomenological analysis of Koenderink, van Doorn, and Braun (2024b). The present three-dimensional metric field reproduces this non-uniformity without imposing it parametrically.

Finally, we examine a planar cross-section orthogonal to the achromatic axis at mid-height within the RGB cube. This plane contains colors of approximately equal luminance and therefore provides a natural setting for comparison with classical chromatic discrimination studies. Figure 17 shows the resulting section. The left panel displays the full three-dimensional ellipsoids at the sampled locations. The right panel shows their intersections with the plane, producing planar ellipses that can be compared directly within a common geometric framework.

**Figure 17.**
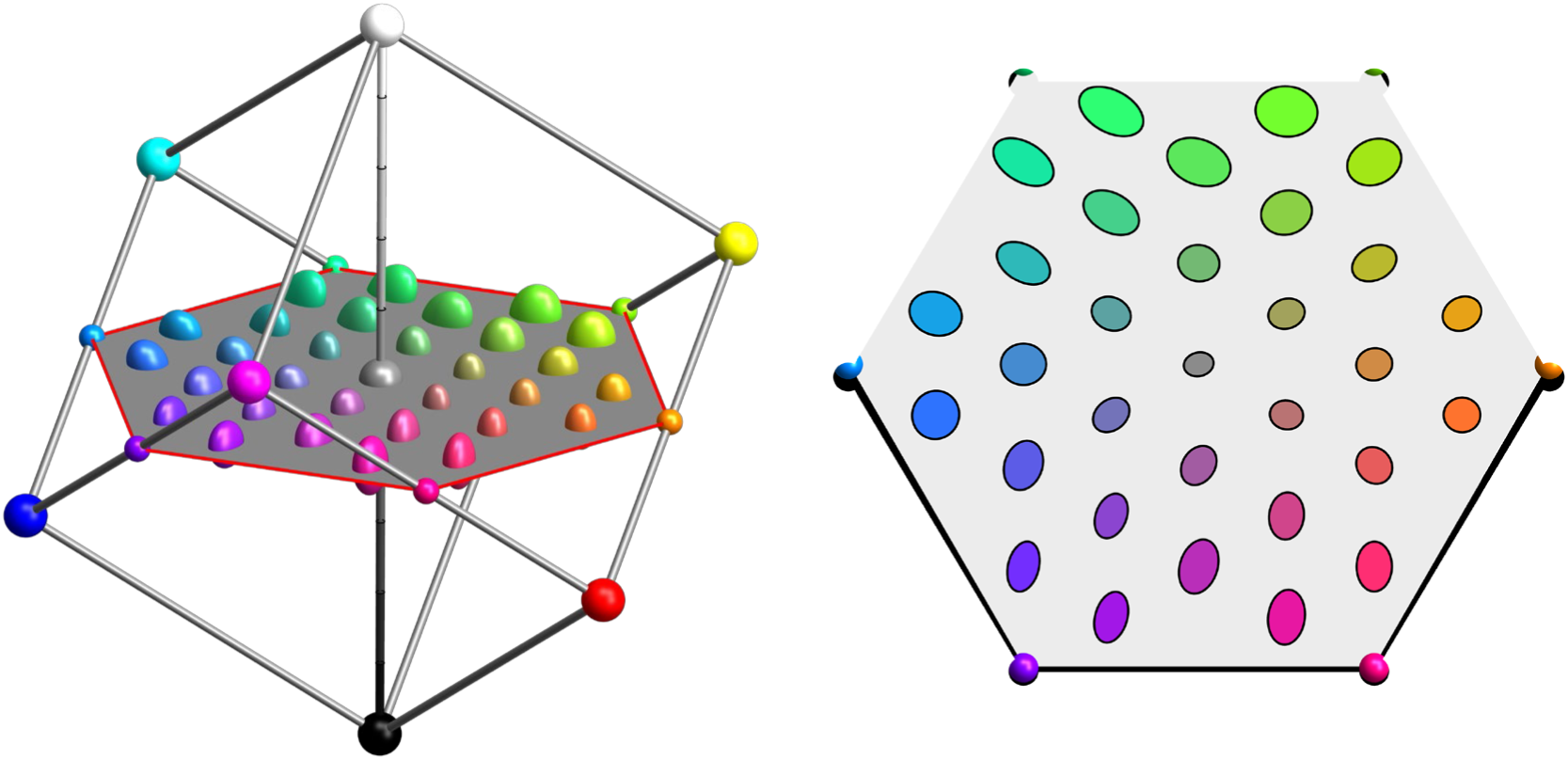
Planar cross-section of the RGB cube at mid-height, orthogonal to the achromatic axis. Left panel shows the full three-dimensional ellipsoids. Right panel shows their intersections with the plane, appearing as flat ellipses that can be directly compared.

Restricting the field to this plane reveals structure that is already present in three dimensions but becomes especially transparent in two. The ellipses vary systematically in both size and orientation. Their major axes are not randomly distributed; instead, neighboring ellipses tend to share similar orientations, forming smooth orientation fields across the section. This representation differs fundamentally from the conventional CIE xy chromaticity diagram. Chromaticity diagrams are obtained by projection and therefore distort geometric relations such as distances, areas, and orientations. By contrast, the present section is an actual plane embedded within the three-dimensional metric field. Relative sizes, shapes, orientations, and distances are therefore represented directly within the plane. The hexagonal representation thus provides a natural planar view of chromatic relations at constant intensity. In this planar view, anisotropies that were previously embedded in three dimensions become directly visible as oriented ellipses. The structure resembles classical MacAdam ellipses in appearance, yet it arises here as a slice through a continuous volumetric metric field rather than as isolated planar measurements.

Our evaluation of the continuous metric field along selected submanifolds reveals a coherent and structured geometry. Along the achromatic axis, grain increases monotonically from black to white. Along chromatic opponent diagonals, grain exhibits V-shaped profiles with systematic asymmetries between complementary directions. Along an interior chromatic circle, qualitative steps vary substantially with hue, confirming that chromatic space is not uniform. A planar section at constant intensity reveals organized anisotropies whose orientations cluster along a small number of dominant chromatic directions, closely resembling the principal axes of variation found in natural color distributions (Koenderink & van Doorn, 2017).

These examples demonstrate that the global metric field is a usable geometric object. It allows structured analysis of one-dimensional subspaces such as the grayscale axis, closed chromatic trajectories such as color circles, planar sections of RGB space, and arbitrary dense samplings throughout the cube. In all cases, the field exhibits smooth and systematic variation. No local irregularities dominate the structure, and the qualitative anisotropies observed earlier are consistently expressed across submanifolds. The metric field thus functions as a coherent geometric description of qualitative color grain. Although visualized here in sRGB coordinates, the same empirical structure can be transferred to linear RGB, XYZ, LMS, DKL, and other color-space representations (Appendix A3, A4).

Having established this empirical geometry, we now compare it to the current operational standard for color difference, CIEDE2000.

### Comparison with the CIEDE2000 Metric

To evaluate how the empirically derived metric field relates to the current operational standard, we constructed a CIEDE2000-derived grain field over the same RGB sample locations and normalized it to the same global scale as the empirical field. The details of this construction are given in Appendix A4. This allows direct geometric comparison independent of overall magnitude. Figure 18 shows the empirical ellipsoids (top row) and the CIEDE2000-derived ellipsoids (bottom row) at the 35 sampled locations. Both fields are displayed at double size and share a common normalization. Several similarities are immediately apparent. The large-scale spatial organization of grain volume is broadly consistent. Regions that are large in the empirical field tend also to be large in the CIEDE2000 field, and small regions correspond similarly. However, systematic differences are visible in both size and orientation. In particular, certain chromatic regions exhibit stronger anisotropy under CIEDE2000, and some orientations differ subtly but consistently. These differences are not random but structured across space.

**Figure 18.**
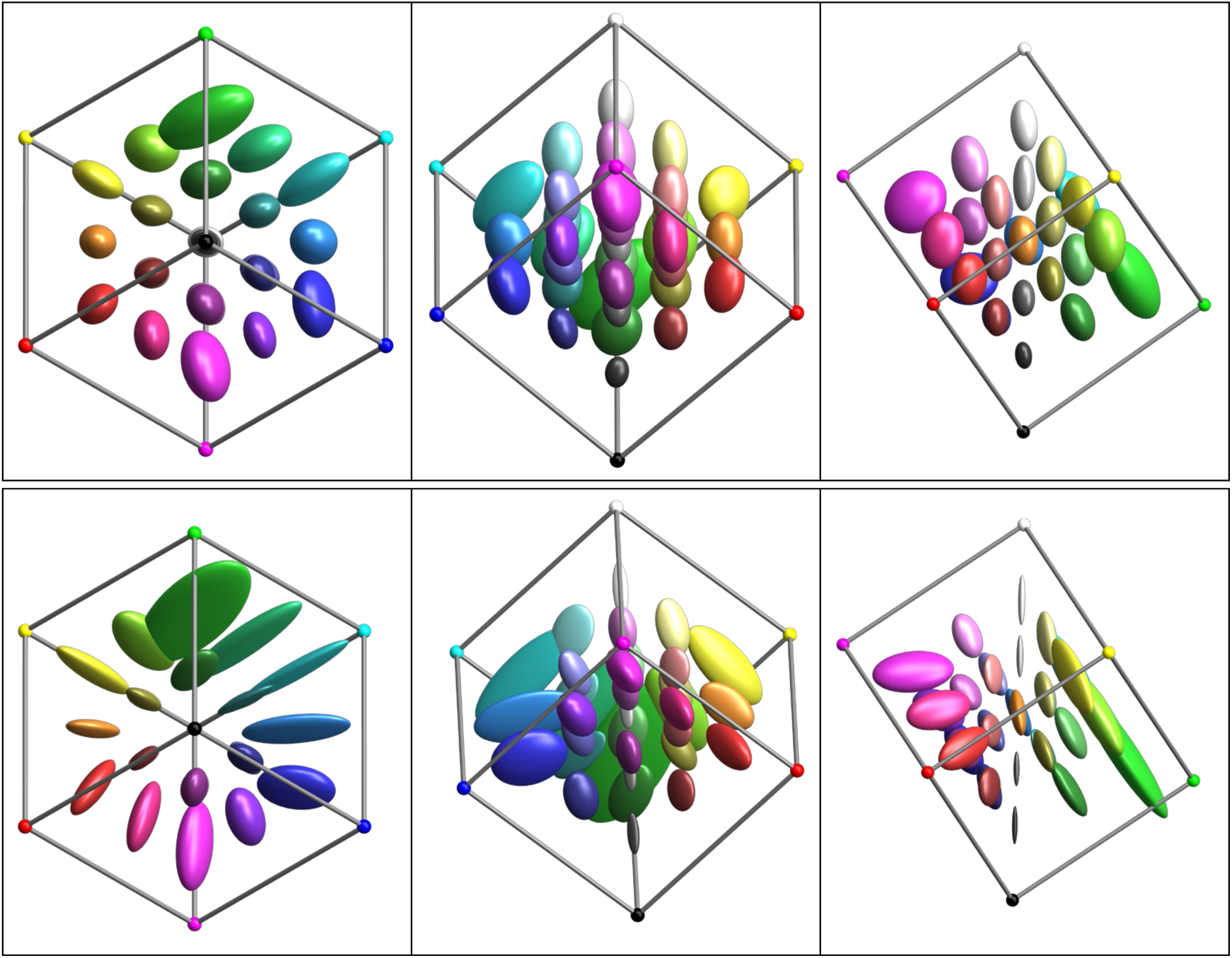
Direct comparison of the empirical grain field (top row) and the CIEDE2000-derived field (bottom row). Ellipsoids are shown at double size and both fields are normalized to the same global scale. Sample points correspond to the measured RGB locations. Broad similarities in spatial structure are evident, alongside systematic differences in anisotropy and orientation.

To quantify these impressions, we compared each individual observer’s ellipsoids to two fields, the empirically derived median field and the CIEDE2000 field. For the median field we used a leave-one-out procedure: each observer was compared against the median of the remaining seven observers, so that no observer contributed to the field it was tested against and the comparison is genuinely out-of-sample. We decomposed the agreement into overall mismatch, volume, elongation and orientation, shown in Figure 19.

**Figure 19.**
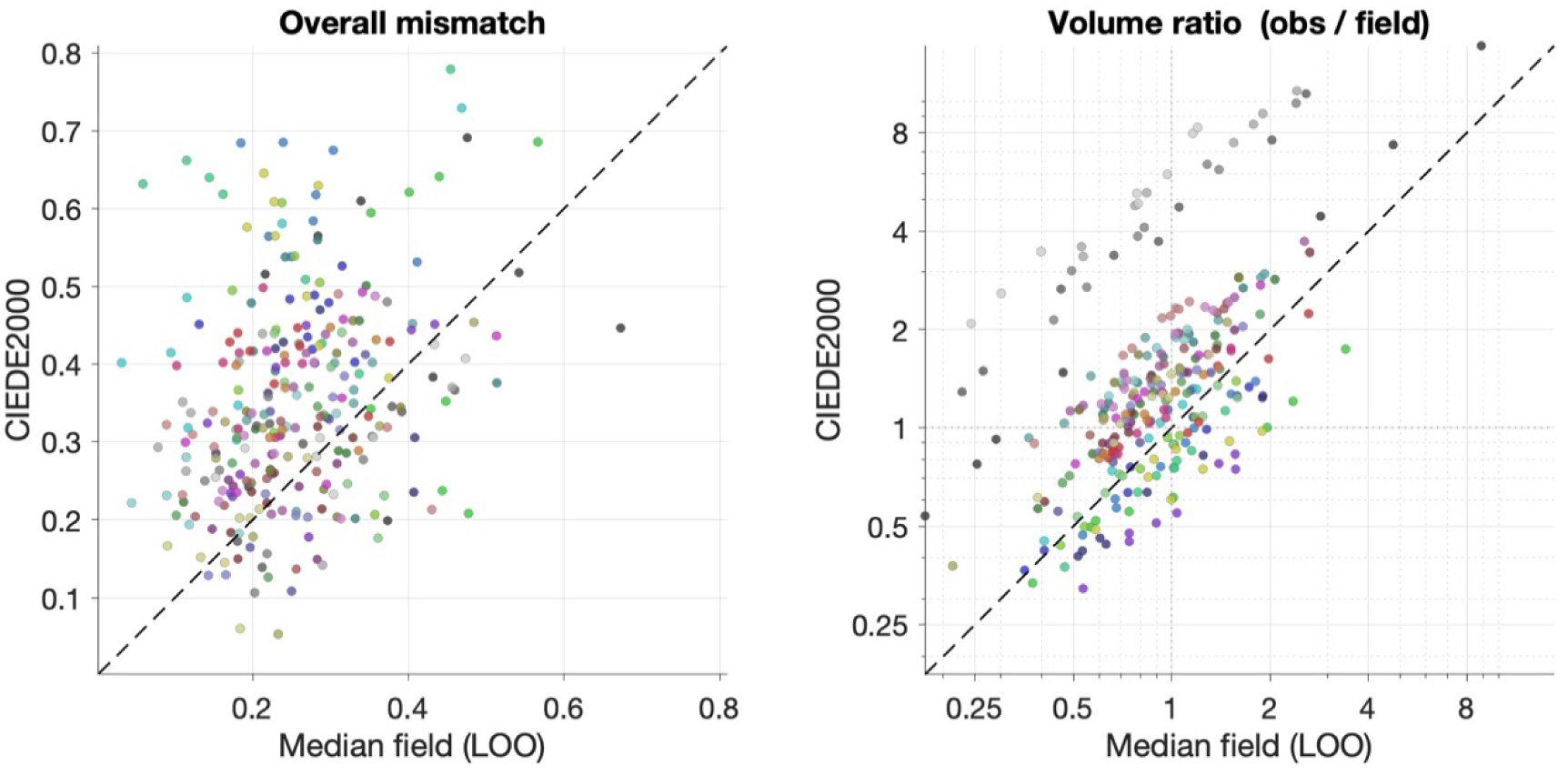
Per-observer goodness of fit of the empirical median field versus CIEDE2000. Each point is one observer × location cell (8 observers × 35 reference colors = 280 points), colored by the location’s RGB. The horizontal axis is the fit to the data-derived median field, evaluated leave-one-out (the median of the other seven observers); the vertical axis is the fit to CIEDE2000 at its single best global scale. The dashed line is equality. Left: total normalized Frobenius mismatch between the observer’s ellipsoid and each field. Right: ratio of ellipsoid volumes on logarithmic axes (gridlines at 1 mark equal size).

The overall mismatch is consistently higher for CIEDE2000 than for the median field (median 0.34 versus 0.24), so on balance the empirical field remains the closer description of the individual observers. Volume, however, agrees well, confirming the qualitative impression from Figure 18. The two fields are strongly rank-correlated in volume across the 35 locations (Spearman’s ρ = 0.91), as high as the agreement between an individual observer and the median field (median ρ = 0.90); CIEDE2000 thus predicts which regions are large or small as reliably as a typical observer does, and the volume ratios cluster around unity. The one clear exception is the achromatic ellipsoids, which CIEDE2000 makes markedly smaller: after matching overall scale, the empirical grayscale ellipsoids are roughly three times larger than their CIEDE2000 counterparts, the discrepancy growing from near-black toward light gray. This reflects the structured monotonic gradient along the grayscale axis noted earlier.

The discrepancy is therefore carried mainly by shape rather than size. CIEDE2000 ellipsoids are systematically more elongated (observer/field elongation ratio 0.65, versus 1.20 for the median field, where values below one indicate a field more elongated than the observer), and their major axes are less well aligned with the observers’ (median major-axis difference 27.5° versus 22.4°). This is the quantitative counterpart of the stronger, structured anisotropy visible in the bottom row of Figure 18. Because it is constructed as a modification of CIELAB with separable lightness, chroma, and hue weightings, it imposes systematic anisotropies that need not coincide exactly with those emerging from volumetric empirical measurement.

Across all four measures the median field is the better fit, and the advantage is statistically reliable. Paired t-tests over the eight observers give a significantly smaller discrepancy for the median field on overall mismatch (t(7) = −6.9, p = 2×10⁻⁴), volume (t(7) = −2.7, p = 0.03), elongation (t(7) = −4.7, p = 0.002) and orientation (t(7) = −3.4, p = 0.01). The advantage holds for every observer individually: the ratio of CIEDE2000 to median-field mismatch ranges from 1.15 to 1.68 across the eight observers, with none favoring CIEDE2000. Thus, while CIEDE2000 captures the broad spatial organization of grain volume, it departs systematically from the observers in shape, and the empirically derived field provides a better account of every individual.

## Conclusions

We measured local regions of notable qualitative color difference at 35 locations throughout the RGB cube and showed that these regions are well approximated by fuzzy ellipsoidal volumes. After normalization for individual scale, observers exhibited a highly consistent spatial structure of qualitative grain. Representing these ellipsoids as symmetric positive-definite matrices enabled averaging and interpolation, yielding a continuous perceptual metric field over RGB space.

Our work suggests a shift in how perceptual color geometry is conceptualized. Rather than seeking a single global distance formula that approximates local discrimination data, we treat color space as endowed with a smoothly varying metric field. This perspective moves the problem from parametric correction to empirical geometry. It replaces adjustments layered onto a predefined coordinate system with a directly measured volumetric structure. The resulting field shows that qualitative color space is neither uniform nor chaotic. It is structured, anisotropic, and smoothly organized across RGB space. Crucially, this structure emerges from finite perceptual steps rather than thresholds. Threshold measurements probe sensitivity at the limit. Qualitative grain probes the scale at which color space is experienced as segmented into distinct regions. These are different levels of description, and they might yield different geometries.

The estimate that RGB space supports about a thousand qualitative regions should not be interpreted as a literal count of color names or a strict capacity limit. It is a geometric statement about graininess. Just as a surface may appear smooth at one scale and granular at another, perceptual color space appears continuous at threshold scale yet discretized at the level of qualitative steps. The coexistence of these scales may be fundamental to how color is represented neurally and cognitively (Griffin & Mylonas, 2019).

The metric field provides a new empirical substrate for theory. Perceptual and neural models of color processing often assume opponent axes, separable dimensions, or low-dimensional embeddings. A volumetric metric offers a quantitative object against which such assumptions can be evaluated. Computational models trained on natural image statistics may now be tested not only on discrimination thresholds but on the structure of qualitative grain.

The present sampling was deliberately sparse yet sufficient to reveal large-scale organization. Future work could refine local structure and boundary regions through denser measurement. High-throughput and immersive paradigms, including gamified approaches such as those recently proposed by Agosti, Hadnett-Hunter, and Gegenfurtner (2026), may make it feasible to collect vastly larger datasets while preserving ecological validity. Such methods could bridge classical threshold psychophysics and volumetric metric reconstruction.

Parametric standards such as CIEDE2000 can be understood as approximations to an underlying perceptual geometry. Direct empirical measurement of that geometry provides a more fundamental reference from which simplified metrics may be derived, tested, or revised. To some extent, larger color differences could be obtained by integrating the local metric along geodesic paths through the field, providing a principled basis for constructing simplified coordinates or distance formulae.

Color science began with the recognition that color matching implies three-dimensional structure. The next step may be to accept that this three-dimensional space is not merely a vector space but a curved and structured manifold endowed with its own intrinsic metric. The present study provides a first empirical chart of that manifold within RGB space.

## Acknowledgments

This work was supported by European Research Council ERC AdG Color 3.0 (884116) and by DFG Sonderforschungsbereich SFB TRR 135 project C2 (222641018). JK was supported by a Humboldt Research Award of the Alexander von Humboldt Foundation.

## Appendix A1: Gamma

Nearly all displays put the digital output values through a non-linear transformation (for review, see Poynton, 2012). This gamma-transformation, named after the exponent of the non-linearity *v* = *x^γ^* was introduced to display technology to counter the effects of Weber’s Law in perception. With increasing luminance, thresholds for luminance differences increase linearly. The gamma transformation makes differences at low intensities smaller and those at high intensities bigger. In vision science, the non-linearity is usually avoided by gamma-correcting output values to achieve a linear relationship between input and output. This makes it possible to relate the input values directly to physiological processes such as cone absorbtions. In our work, we are faced with two questions: First, is there any harm done by not gamma-correcting our display. Second, are there possibly any advantages to doing so.

The transformation from gamma-corrected sRGB to linear RGB (and hence to XYZ or LMS) is a known, monotonic, smooth function. The NQD regions remain NQD regions after transformation, only their shape in the new coordinate system changes. The metric field can therefore be evaluated in any target space by transforming the ellipsoid matrices at each reference location. We performed the complete analysis also on linear RGB Values and compared the result to the version of the metric field that was transformed from non-linear to linear RGB by analytical methods. As we will see below, the results are virtually identical and the result can be used to go directly to any other linearly related color space such as LMS, XYZ or DKL.

We would like to point out, however, that there is a distinct advantage to using non-linear RGB. The obvious point is that it is most directly related to what is being displayed, and that it’s the color space most people work in. The more subtle point is that non-linear RGB is an approximately whitened, stationary representation of the perceptual metric, for the same reason L* is. The non-linearity defining the L* intensity coordinate of CIELAB color space, is essentially identical to the gamma transform applied in displays. What is good for the most common CIE-defined color space couldn’t be that bad for RGB then.

**Figure A1.**
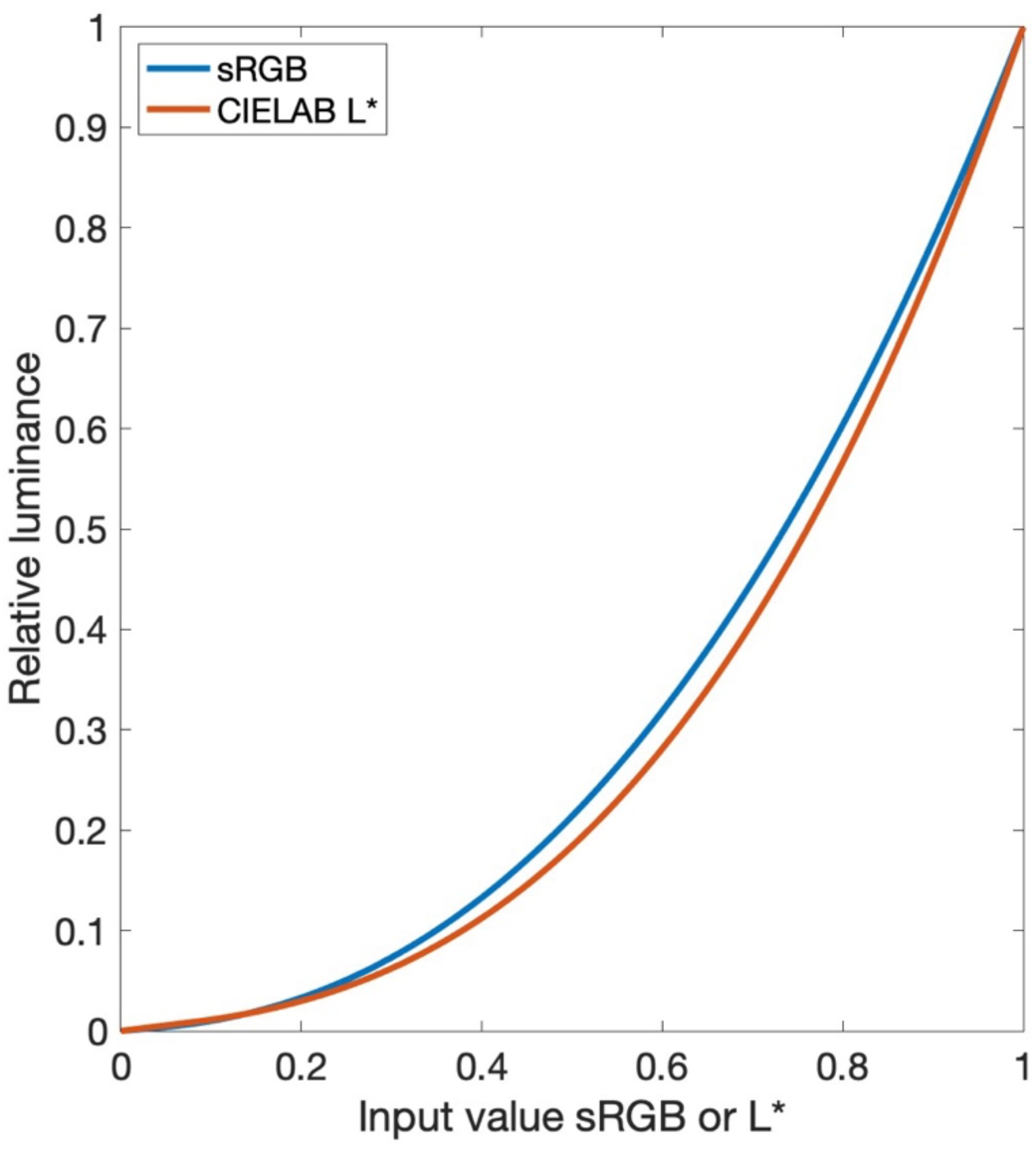
Comparison of nonlinear intensity encodings. The sRGB gamma function (γ≈2.4) and the cube-root nonlinearity underlying the CIELAB lightness specification produce very similar compressions of the physical intensity scale over most of the range relevant for display stimuli. The difference between the two functions is small compared with the variability of the perceptual grain measurements reported here. L* is scaled between 0 and 1 in this graph.

The major advantage for our measurements is that most ellipsoids won’t be characterized by a massive elongation along the intensity axis. They will be more stationary, and deviations from the intensity axis will be easier to measure reliably. It should be noted that this approach is widely used in other domains of sensory science. Sound pressure in pascals follows linear superposition (the real world is linear), yet every acoustician works in decibels without apology, because dB is what makes the perceptual metric stationary. We just apply this approach to color.

## Appendix A2: The empirical median metric field

**Table A2:**
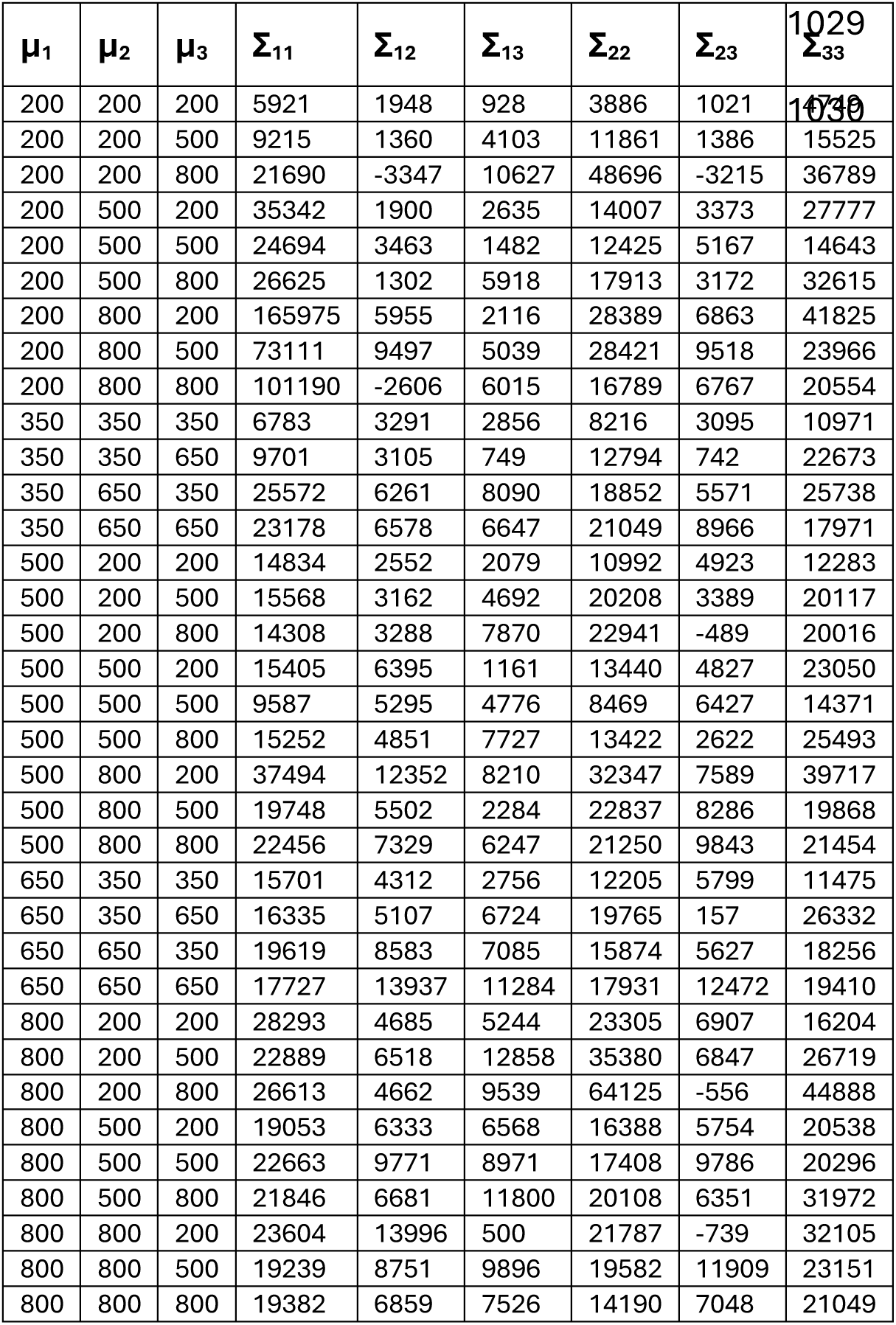
RGB locations and Sigma matrices. Locations μ₁–μ₃ are ×1000. Matrix coefficients Σᵢⱼ are ×10,000,000.

## Appendix A3: The linear metric field

The local NQD regions measured in this study are perceptual entities and therefore independent of the coordinate system used to represent them. Any smooth one-to-one transformation of color space maps NQD regions into corresponding NQD regions in the transformed space. Only their geometric representation changes.

Consequently, the entire analysis presented in this paper could, in principle, be repeated in any desired color space. For a target space such as linear RGB, the staircase endpoints may first be transformed from gamma-encoded sRGB coordinates. Convex hulls can then be computed, minimum-volume circumscribed ellipsoids fitted, observer medians calculated, and the metric field reconstructed exactly as described for the original data. We refer to this procedure as **Method A**.

The resulting linear-RGB metric field has a special status because linear RGB is related to XYZ, LMS, and DKL coordinates by linear transformations. Once a metric field has been constructed in linear RGB, ellipsoids may be transferred to any linearly related space without further fitting. If

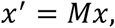

then an ellipsoid represented by

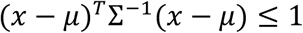

is transformed according to

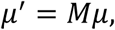

and

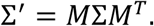

Thus a single metric field in linear RGB is sufficient to generate corresponding fields in XYZ, LMS, DKL, or any other linearly related representation.

A practical complication arises because the transformation from gamma-encoded sRGB to linear RGB is nonlinear. An ellipsoid in gamma space therefore does not generally transform into an exact ellipsoid in linear RGB. In principle, one should transform the original NQD regions and determine their geometry anew. Fortunately, the deviations from ellipsoidal symmetry in the original data are already modest, with median radial asymmetries of approximately 10%. After transformation to linear RGB, the asymmetries remain of the same magnitude. The ellipsoidal approximation therefore remains equally appropriate in the transformed space.

To evaluate the consequences of the nonlinear transformation, we compared three approaches for constructing the metric field in linear RGB.

**Method A** transformed the original staircase endpoints into linear RGB and repeated the complete analysis. Convex hulls were computed from the transformed endpoints, minimum-volume circumscribed ellipsoids were fitted, observer medians were calculated, and the metric field was reconstructed exactly as for the gamma-encoded data. This procedure serves as the reference method because it starts from the original psychophysical measurements.

**Method B** started from the fitted gamma-space ellipsoids. Each ellipsoid was densely sampled on its surface, the 300 sample points were transformed into linear RGB, and a new ellipsoid was fitted to the transformed point cloud. This method approximates the exact transformation of the ellipsoidal regions themselves and avoids repeating the entire analysis pipeline.

**Method C** used a local analytical approximation. The transformed ellipsoid was obtained from the Jacobian of the sRGB-to-linear transformation evaluated at the ellipsoid center,

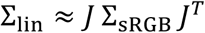

Because the transformation acts independently on the three color channels, the Jacobian is diagonal and the computation is straightforward.

Agreement between the three methods was quantified using the normalized Frobenius mismatch. Across all observers and sampled locations, the median mismatch between the direct linear fit (Method A) and the geometric transformation (Method B) was 0.041. The mismatch between Method A and the Jacobian approximation (Method C) was 0.050. The mismatch between the two transformation approaches (Methods B and C) was only 0.021. Figure 7 of the main paper illustrates that deviations of this magnitude are negligible.

The same conclusion holds for the observer-aggregated median field. Across the 35 sampled locations, median mismatches were 0.042 for Method A versus Method B, 0.044 for Method A versus Method C, and 0.006 for Method B versus Method C.

The conclusion is straightforward. The geometric structure of the metric field is largely invariant to whether it is estimated directly in gamma-encoded sRGB coordinates or reconstructed in linear RGB space^2^. The coordinate representation affects local ellipsoid geometry only minimally relative to the intrinsic variability of the measurements. A metric field constructed in linear RGB may therefore be transformed with confidence into XYZ, LMS, DKL, and related color spaces using the corresponding linear coordinate transformations.

**Figure A3.**
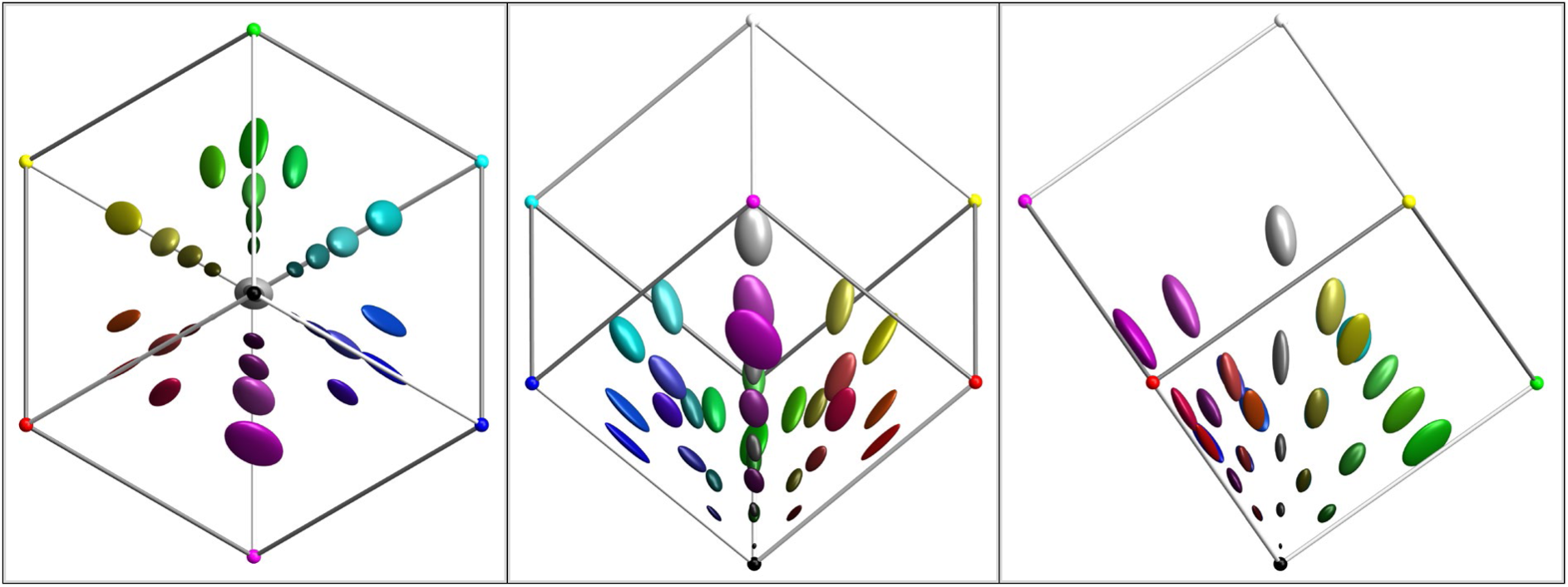
The equivalent of Figure 8 in linear RGB space. Three orthogonal views are provided. Left panel is viewed along the white–black axis, center panel along the purple–green axis, and right panel along the orange–teal axis. The field varies smoothly in both size and orientation across linear RGB space

**Table A3:** Linear RGB locations and Sigma matrices.

| $\mu_1$ | $\mu_2$ | $\mu_3$ | $\Sigma_{11}$ | $\Sigma_{12}$ | $\Sigma_{13}$ | $\Sigma_{22}$ | $\Sigma_{23}$ | $\Sigma_{33}$ |
| --- | --- | --- | --- | --- | --- | --- | --- | --- |
| 33 | 33 | 33 | 424 | 128 | 57 | 269 | 71 | 334 |
| 33 | 33 | 214 | 653 | 108 | 843 | 807 | 315 | 8739 |
| 33 | 33 | 604 | 1554 | -108 | 3913 | 3552 | -754 | 61693 |
| 33 | 214 | 33 | 2779 | 515 | 608 | 6615 | 924 | 1867 |
| 33 | 214 | 214 | 2030 | 872 | 371 | 6947 | 2721 | 7568 |
| 33 | 214 | 604 | 2055 | 435 | 2461 | 10024 | 3732 | 59316 |
| 33 | 604 | 33 | 23168 | 4171 | 1538 | 51057 | 3783 | 3286 |
| 33 | 604 | 214 | 8284 | 4471 | 1708 | 47320 | 9262 | 11640 |
| 33 | 604 | 604 | 9380 | -344 | 3415 | 26785 | 10328 | 34202 |
| 100 | 100 | 100 | 1513 | 758 | 696 | 1780 | 741 | 2526 |
| 100 | 100 | 380 | 2368 | 697 | 428 | 2789 | 507 | 22491 |
| 100 | 380 | 100 | 5928 | 3521 | 2134 | 20862 | 2998 | 5832 |
| 100 | 380 | 380 | 5229 | 3125 | 3724 | 20563 | 10336 | 19013 |
| 214 | 33 | 33 | 7529 | 528 | 452 | 733 | 370 | 764 |
| 214 | 33 | 214 | 7386 | 777 | 2643 | 1506 | 829 | 11390 |
| 214 | 33 | 604 | 8501 | 714 | 7712 | 2009 | 36 | 35963 |
| 214 | 214 | 33 | 7784 | 3848 | 268 | 8011 | 1057 | 1723 |
| 214 | 214 | 214 | 5527 | 2800 | 2536 | 5291 | 3967 | 7675 |
| 214 | 214 | 604 | 8559 | 2656 | 8189 | 7199 | 2796 | 48261 |
| 214 | 604 | 33 | 19619 | 11731 | 2577 | 51236 | 3214 | 3321 |
| 214 | 604 | 214 | 10356 | 5613 | 1516 | 38579 | 9030 | 11466 |
| 214 | 604 | 604 | 11915 | 8138 | 6193 | 36188 | 17974 | 36632 |
| 380 | 100 | 100 | 15835 | 2179 | 1351 | 2668 | 1480 | 3041 |
| 380 | 100 | 380 | 16468 | 3084 | 8276 | 4689 | 464 | 29520 |
| 380 | 380 | 100 | 19876 | 8723 | 3526 | 15945 | 2811 | 4325 |
| 380 | 380 | 380 | 18206 | 13957 | 13147 | 18867 | 13097 | 20811 |
| 604 | 33 | 33 | 44800 | 1764 | 1786 | 1876 | 538 | 1222 |
| 604 | 33 | 214 | 41596 | 2852 | 13244 | 3324 | 1919 | 14431 |
| 604 | 33 | 604 | 42732 | 2645 | 18655 | 6053 | 913 | 81143 |
| 604 | 214 | 33 | 36105 | 7445 | 2622 | 10062 | 1108 | 1499 |
| 604 | 214 | 214 | 39361 | 10384 | 9322 | 9574 | 5384 | 11119 |
| 604 | 214 | 604 | 36936 | 6956 | 23409 | 11485 | 6622 | 60644 |
| 604 | 604 | 33 | 39844 | 23614 | 253 | 36596 | -136 | 2723 |
| 604 | 604 | 214 | 35190 | 16949 | 10066 | 31964 | 11534 | 12360 |
| 604 | 604 | 604 | 32065 | 12629 | 12736 | 24849 | 13058 | 39018 |

## Appendix A4: Construction of DE2000 ellipsoids

The CIEDE2000 color-difference measure is defined as a nonlinear numerical algorithm rather than as a simple analytical expression. Strictly speaking, it is not a distance function in the formal mathematical sense, because it does not fully satisfy the triangle inequality. Nevertheless, it has been widely adopted to describe perceived color differences between similar colors, based on committee agreements that synthesize diverse psychophysical datasets and practical experience in industrial color evaluation.

The CIEDE2000 measure is intended for the comparison of related colors. In the present analysis we restrict the domain to the RGB unit cube 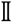^3^ ⊂ ℝ^3^ viewed as a subset of ℝ³, while the range consists of the non-negative real numbers. In abstract terms, the color-difference measure can be regarded as a function

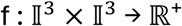

defined on the RGB unit cube 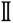^3^ such that *f(k,k)* = 0 and *f(k,l)* > 0 whenever *k* ≠ *l*. For our purposes we seek a local metric representation of this distance function.

Specifically, we determine for each reference color *k* a symmetric positive-definite matrix *Σ(k)* such that

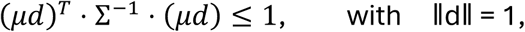

describes a perceptually uniform ellipsoidal region around k. The scalar factor μ may be interpreted as the corresponding Δ*E* value.

Several numerical strategies could be used to determine Σ(k). We chose to probe the neighborhood of each point *k* along a set of uniformly distributed directions. In practice we used 32 directions; increasing this number produced no noticeable change in the resulting ellipsoids.

For each direction *dᵢ* we compute the color difference

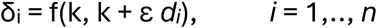

and approximate the directional derivative of the distance function by δᵢ / ε. The increment ε was chosen as 10⁻⁵. Smaller values produced no systematic changes and may introduce numerical inaccuracies due to floating-point precision.

The points *μᵢ dᵢ*, with *μᵢ* = ΔE·ε/δᵢ, lie on the Δ*E* isosurface of the color-difference function. A minimum-volume bounding ellipsoid algorithm is then used to determine the corresponding matrix Σ(k). In practice we employed a Lӧwner–John ellipsoid fit to the sampled points (Güler & Gürtuna, 2012).

This approach has the advantage that it does not rely on explicit differentiation of the CIEDE2000 formula, which is algebraically complex and not specified in closed form. Instead, the method samples the local behavior of the distance function directly and constructs the ellipsoid numerically.

The resulting ellipsoids provide a local ellipsoidal approximation to the ΔE isosurfaces of the CIEDE2000 color-difference function within RGB space.

## Appendix A5: Display characteristics

**Figure A5:**
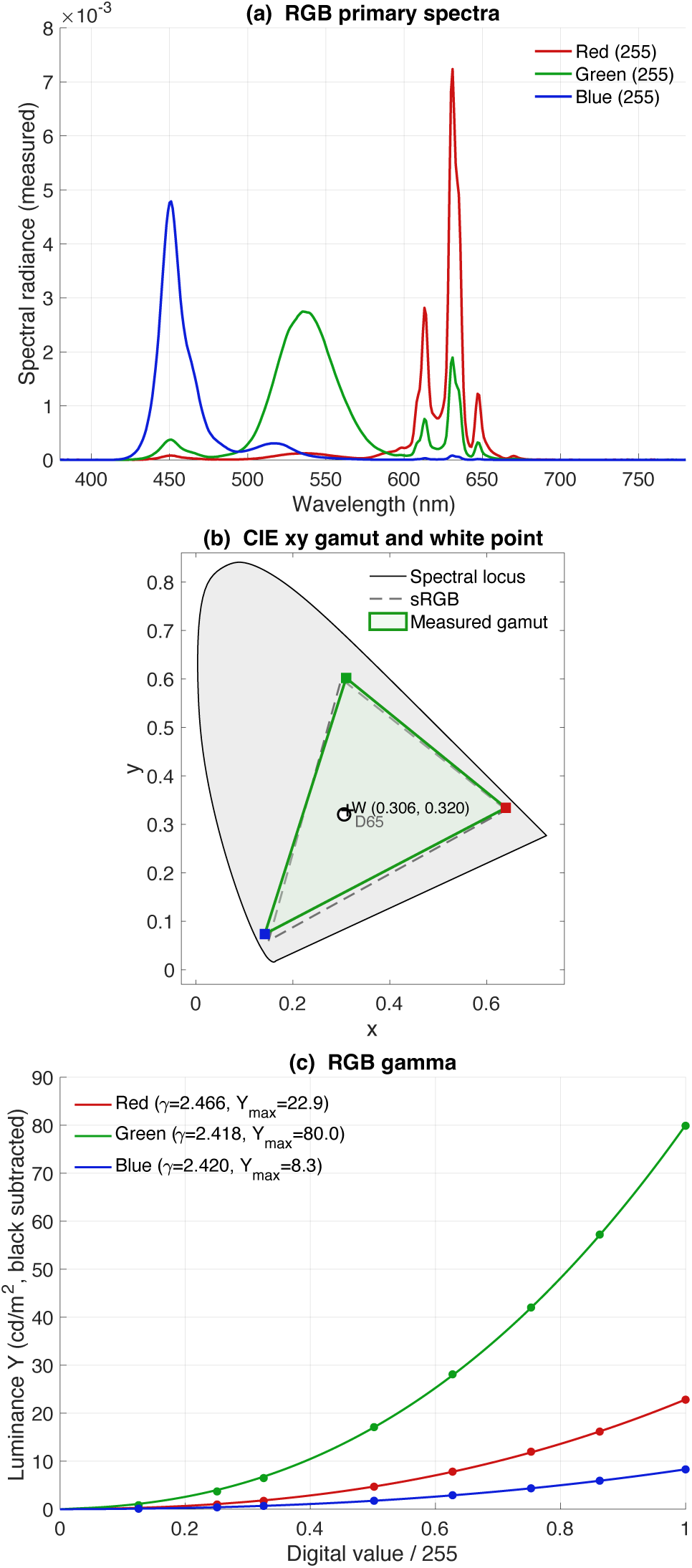
(Top) RGB primary spectra of the Apple MacBook used to run the experiments. (Middle) Display gamut (Bottom) Gamma curves.

**Table A5:** Spectral emission curves of our display primaries.

| nm | R | G | B |
| --- | --- | --- | --- |
| 400 | 0 | 1.094E-06 | 1.1168E-06 |
| 410 | 1.894E-07 | 1.2523E-06 | 2.8821E-07 |
| 420 | 8.8214E-07 | 1.4159E-06 | 1.7991E-05 |
| 430 | 3.9016E-06 | 1.6828E-05 | 0.00020395 |
| 440 | 2.2616E-05 | 0.00011283 | 0.0014257 |
| 450 | 7.9261E-05 | 0.00037158 | 0.004762 |
| 460 | 3.9618E-05 | 0.00017894 | 0.0022927 |
| 470 | 1.7223E-05 | 8.3695E-05 | 0.0010065 |
| 480 | 8.1024E-06 | 5.8154E-05 | 0.00036077 |
| 490 | 7.7396E-06 | 0.00011236 | 0.00017044 |
| 500 | 1.4704E-05 | 0.00029174 | 0.00017602 |
| 510 | 3.9342E-05 | 0.00086919 | 0.00027334 |
| 520 | 8.3275E-05 | 0.0018402 | 0.00030175 |
| 530 | 0.00011503 | 0.0025753 | 0.00017845 |
| 540 | 0.00011918 | 0.0027142 | 7.7345E-05 |
| 550 | 9.3323E-05 | 0.0020962 | 4.1282E-05 |
| 560 | 5.5831E-05 | 0.0012764 | 2.2477E-05 |
| 570 | 3.0997E-05 | 0.00064454 | 1.1961E-05 |
| 580 | 5.7445E-05 | 0.00027109 | 5.1155E-06 |
| 590 | 0.00014142 | 0.00011809 | 3.1573E-06 |
| 600 | 0.00021994 | 8.1017E-05 | 2.7815E-06 |
| 610 | 0.0013115 | 0.00036343 | 1.4366E-05 |
| 620 | 0.00079709 | 0.00021033 | 8.8148E-06 |
| 630 | 0.0068261 | 0.0017877 | 7.4939E-05 |
| 640 | 0.00046164 | 0.00012196 | 5.6506E-06 |
| 650 | 0.00058739 | 0.00015358 | 7.4642E-06 |
| 660 | 8.2411E-05 | 2.2487E-05 | 1.2275E-06 |
| 670 | 7.2962E-05 | 1.9287E-05 | 7.2229E-07 |
| 680 | 7.7471E-06 | 2.1165E-06 | 1.8781E-07 |
| 690 | 4.9629E-06 | 1.907E-06 | 0 |
| 700 | 2.0911E-06 | 2.7128E-07 | 0 |

## Footnotes

1 For an animated display of the metric field, see https://www.allpsych.uni-giessen.de/karl/ellie/

2 For an animated display of the metric field, see https://www.allpsych.uni-giessen.de/karl/ellie, where you can switch between linear and gamma-encoded versions.

